# A Commensal-Derived Lipoteichoic Acid Engages an Inducible Neuronal PD-1 Checkpoint to Suppress Inflammatory Pain

**DOI:** 10.64898/2026.01.16.699990

**Authors:** Zhiqiang Liu, Yen-Hsin Cheng, Catherine V. Osborn, Marco Martina, Anthony J. Schaeffer, Praveen Thumbikat

**Affiliations:** Department of Urology, Northwestern University Feinberg School of Medicine, Chicago, IL, United States; Department of Neuroscience, Northwestern University Feinberg School of Medicine, Chicago, IL, United States

**Keywords:** PD-1, sensory neurons, analgesia, *Staphylococcus epidermidis*, lipoteichoic acid, SELTA, neuroimmune modulation, chronic pain, inflammatory pain

## Abstract

**Background:** Immune checkpoint receptors regulate adaptive immunity but are increasingly recognized as modulators of neuroimmune interactions. The upstream signals that induce neuronal checkpoint pathways during inflammation and their functional relevance in inflammatory pain remain incompletely understood. We investigated whether a defined commensal-derived molecule engages a neuroimmune checkpoint axis to modulate inflammatory pain.

**Methods:** SELTA, a lipoteichoic acid purified from a commensal *Staphylococcus epidermidis* strain, was evaluated for TLR2-dependent activity, regulation of Pdcd1 transcription and PD-1 protein expression in dorsal root ganglion (DRG) neurons, effects on intracellular calcium signaling, and behavioral outcomes in experimental autoimmune prostatitis (EAP) a model of inflammation-induced chronic pelvic pain. Conditional Pdcd1 deletion was performed in sensory neurons (Advillin-^Cre^) and CD4⁺ T cells to determine cell-specific requirements.

**Results:** SELTA selectively activated TLR2/6 signaling and increased Pdcd1 transcription and PD-1 protein expression in DRG neurons under inflammatory conditions. SELTA enhanced phosphorylation of PD-1 at tyrosine 248 and significantly reduced ATP-evoked intracellular Ca²⁺ responses in mouse primary sensory neurons. Pharmacologic neutralization of PD-1 abrogated SELTA-mediated suppression of calcium signaling. In vivo, SELTA produced concentration-dependent attenuation of pelvic hypersensitivity in EAP. Conditional deletion of PD-1 *in vivo* in Advillin-expressing sensory neurons or CD4⁺ T cells significantly reduced SELTA-induced analgesia, while combined deletion did not further diminish the effect.

**Conclusions:** These findings identify a commensal-derived lipoteichoic acid that engages a TLR2/6-associated pathway linked to inducible neuronal PD-1 signaling to restrain inflammatory nociceptor activity. The results define a neuroimmune checkpoint mechanism that modulates inflammatory pain and extend PD-1 signaling beyond its established role in adaptive immunity.

## Introduction

Chronic pain affects more than 20% of adults worldwide and is a leading cause of disability and opioid exposure [7]. Chronic pelvic pain syndromes, including chronic prostatitis/chronic pelvic pain syndrome (CP/CPPS), are characterized by persistent peripheral sensitization and nociceptor hyperexcitability yet remain difficult to treat effectively [4; 5; 32]. Although pro-inflammatory mediators contributing to pelvic pain have been extensively characterized [18], endogenous inhibitory pathways that limit sensory neuron activity during chronic inflammation are less well defined. Identification of such mechanisms is critical for the development of non-opioid therapeutic strategies.

The immune and nervous systems share bidirectional regulatory circuits that coordinate host defense and tissue homeostasis. Recent work has demonstrated that the immune checkpoint receptor programmed cell death-1 (PD-1), classically known for limiting T-cell activation, is also expressed in dorsal root ganglion (DRG) neurons where it suppresses neuronal excitability and inflammatory pain [3; 34]. Engagement of PD-1 by its endogenous ligands PD-L1 or PD-L2 reduces nociceptor firing, whereas PD-1 blockade enhances pain sensitivity and interferes with opioid antinociception in preclinical models and clinical settings [3; 8; 33]. These findings suggest that PD-1 functions as a neuroimmune checkpoint regulating peripheral sensitization. However, the physiological contexts in which neuronal PD-1 is engaged and whether these exogenous signals can activate this pathway are incompletely understood.

Commensal microbes produce bioactive molecules capable of shaping epithelial and immune signaling [2; 19]. Our laboratory previously identified a lipoteichoic acid (LTA) molecule purified from a commensal *Staphylococcus epidermidis* strain that produces robust analgesia in murine models of pelvic pain [15–17]. This molecule, termed SELTA, showed context-dependent immunomodulatory properties, with minimal activity in quiescent tissue but measurable effects in inflammatory settings [17]. Lipoteichoic acids are recognized ligands of Toll-like receptor 2 (TLR2), particularly TLR2/6 heterodimers [27]. Whether TLR2-dependent signaling in sensory neurons can induce inhibitory checkpoint pathways has not been examined.

Baseline single-cell RNA sequencing of naïve DRG demonstrates low expression of both Tlr2 and Pdcd1 transcripts [30], suggesting that checkpoint engagement may be inducible rather than constitutive in sensory neurons. Given the established inhibitory role of PD-1 in nociceptors and the context-dependent activity of SELTA, we hypothesized that SELTA activates TLR2/6-dependent signaling and induces inflammation-dependent PD-1 expression and phosphorylation in sensory neurons. Here, we demonstrate that SELTA utilizes TLR2/6 to activate NF-κB signaling, increases Pdcd1 transcription in DRG resulting in PD-1 phosphorylation under inflammatory conditions, suppresses ATP-evoked calcium signaling, and reduces pelvic hypersensitivity in experimental autoimmune prostatitis (EAP). Using Advillin^cre^-mediated deletion of Pdcd1 in primary sensory neurons and CD4-mediated deletion in T cells, we further show that SELTA-induced analgesia requires PD-1 expression in both neuronal and adaptive immune compartments.

## Materials and Methods

### Ethics statement

All animal procedures were approved by the Institutional Animal Care and Use Committee (IACUC) at Northwestern University (protocols IS00021559 and IS00003610). The IACUC at Northwestern is AAALAC accredited, and Northwestern University has an Animal Welfare Assurance on file with the Office of Laboratory Animal Welfare (A3283-01). All studies were conducted in accordance with United States Public Health Service regulations, the Animal Welfare Act, and applicable federal and local laws.

### Animals

All procedures were approved and monitored by the Northwestern University IACUC. Pain-related experiments were conducted in accordance with the IASP Guidelines for the Use of Animals in Research (January 2014). Male C57BL/6 (B6) and conditional knockout (CKO) mice (6–8 weeks old; Jackson Laboratory, Bar Harbor, ME) were housed with ad libitum access to sterilized food and water. Unless otherwise indicated, 4–5 mice were used per group, and experiments were performed in duplicate or triplicate (N = 2–3) with no exclusion of animals.

### SELTA extraction and purification

SELTA used for experiments (100 ng/ml) was extracted from a unique strain of *S. epidermidis* isolated from the prostatic secretion of a healthy human volunteer at Northwestern University [17] and purified at the Complex Carbohydrate Research Center (CCRC) at the University of Georgia. Briefly, extraction was performed from pelleted bacterial cells suspended in 100 mM Na citrate, pH 4.7 by pipetting. 1-butanol was added and the mixture was shaken vigorously (280 rpm) for 30 min at room temperature. The suspension was repeatedly centrifuged (3X) at 13000g, and the bottom yellow water-rich phase was removed was dialyzed in 1 kDa Mw cut-off bag against 20 mM Na citrate, pH 4.7 overnight at 4 °C and the retentate (∼400 mL) was lyophilized. Purification was performed using hydrophobic interaction chromatography (HIC) chromatography using an HPLC system (Agilent 1260 Infinity II). The material was eluted at 0.3 mL/min using 200 mL linear gradient from buffer A to [41 mM Na citrate, pH 4.7, 65 vol% 1-propanol], and 5 mL fractions were collected for analysis. Following purification, total carbohydrate and phosphorus content was assayed using standard phenol-sulphuric-acid and total phosphorus assays. Finally, NMR spectroscopy was performed to confirm structural information. For this, an aliquot of the purified SELTA was dried on a speedvac, dissolved in 510 μL D2O (99.9% D) and transferred into a 5 mm NMR tube. NMR data were collected at 25 °C on a Bruker Avance III or Varian VNMRS spectrometer (1H, 600 MHz), each equipped with a cryogenically cooled probe. 1H data were acquired with a spectral width of 9615 Hz, 32k complex data points and 64 transients. Prior to the Fourier transformation, NMR data were apodized with an exponentially decaying function (lb = 0.3 Hz) and baselines of the spectra were corrected automatically using an 3rd-order polynomial. The spectra were referenced to citrate signal at 3.681 ppm. Based on integration of NMR signals, the molecular weight of purified deacylated SELTA was determined to be 8.1kDa. Multiple lots of SELTA have been tested to confirm lot-level reproducibility and reduce batch-specific concerns [17]. For *in vivo* treatment, 10µl of the SELTA diluted in water (or saline in controls) was administered to anesthetized male mice through a catheter introduced into the urethra [17]. Mice were maintained under anesthesia for 15-30 minutes followed by recovery and transfer to housing.

### Induction of experimental autoimmune prostatitis (EAP)

EAP was induced in C57BL/6 mice by subcutaneous injection behind the shoulder of a 1:1 emulsion of rat prostate homogenate and TiterMax® adjuvant, as described previously [21]. Pelvic tactile allodynia was assessed using von Frey filament testing. Unless otherwise specified, testing was performed at baseline (prior to EAP induction) and at 7-day intervals thereafter. Experimental endpoints were at day 28 or day 120 post-EAP induction. In SELTA experiments, therapeutic administration was performed one day prior to efficacy testing, following collection of baseline (pre-treatment) measurements. Results were expressed as percentage change relative to baseline, and for each von Frey filament, responses were averaged within each group. Behavioral testing was performed by an experimenter blinded to treatment allocation. At designated endpoints, mice were euthanized and lumbar and sacral DRG were harvested for analysis.

### Generation of cell type–specific conditional knockout mice

*<u>Sensory neuron**–** specific Pdcd1 conditional knockout mice</u>* were generated using a Cre–loxP strategy. Pdcd1 mice harboring loxP sites flanking critical exons of the Pdcd1 gene (gift from Dr. Vassiliki A. Boussiotis) [1] were crossed with Advillin^Cre^ transgenic mice (Jackson Laboratory, stock #032536), which express Cre recombinase under the sensory neuron–specific Advillin promoter. Offspring were genotyped to identify mice homozygous for the floxed Pdcd1 allele and heterozygous for Cre (*Pdcd1* Advillin) (Supplementary Fig. S5). Genomic DNA was extracted from tail biopsies. PCR was performed using the following primers: Pdcd1 floxed allele: Forward: 5′-TATCCCTGTATTGCTGCTGCTG-3′ Reverse:5′-AATGAATTGAGGAGTAGGGCCTG-3′ (387 bp wild-type; 490 bp floxed). Cre transgene: Forward: 5′-AATGGCTCCCTGTTCACTGT-3′ Reverse WT: 5′-TGACTAGGTAGAGGTGCAAATGTC- 3′ (530 bp wild type). Reverse Cre: 5′-AGGCAAATTTTGGTGTACGG-3′ (150 bp transgene). PCR products were resolved on 1.5% agarose gels (Supplementary Fig. S6). For validation, DRG sensory neurons from *Pdcd1^fl/fl^*;Advillin^cre+/−^ and *Pdcd1^fl/f^* littermates were analyzed for PD-1 immunoreactivity by immunohistochemistry to confirm loss of PD-1 in neurons and preservation in non-neuronal cells.

*<u>CD4^+^T lymphocyte–specific Pdcd1 conditional knockout mice</u>* were generated by crossing *Pdcd1*^flox/flox^ mice with B6.Cg-Tg(Cd4-cre)1Cwi/BfluJ mice (The Jackson Laboratory, stock #022071), which express Cre recombinase under the CD4 promoter. Offspring were genotyped to identify *Pdcd1^fl/f^*;CD4^cre+/−^ mice. Genomic DNA was isolated from tail biopsies and PCR amplification was performed using primer sets specific for the CD4^cre^ transgene according to the Jackson Laboratory genotyping protocol for B6.Cg-Tg(Cd4-cre)1Cwi/BfluJ (stock #022071). The transgene-specific reaction yields a 741 bp amplicon. Efficient deletion of PD-1 in CD4⁺ T cells was validated by PCR genotyping, immunofluorescence staining of lymph node sections, and flow cytometric analysis of splenocytes. Immunohistochemistry confirmed loss of PD-1 immunoreactivity in CD4^+^ T cell zones of *Pdcd1^fl/f^* CD4^cre+/−^ mice relative to *Pdcd1* littermate controls, and flow cytometry demonstrated marked reduction of PD-1 expression in CD4⁺ T cells (Supplementary Fig. S6).

*<u>Double conditional knockout mice</u>* lacking PD-1 in both sensory neurons and CD4⁺ T cells, were generated using breeding pairs established with *Pdcd1^fl/f^*;CD4^cre+/−^ females and *Pdcd1^fl/f^*;Advillin^cre+/−^ males. Offspring carrying both Cre alleles (*Pdcd1^fl/f^*/CD4^Cre+/^; Advillin^Cre+/−^) were identified using the genotyping strategies described above and designated as double conditional knockout mice (Supplementary Fig. S6).

### Mechanical allodynia

Mechanical hypersensitivity was assessed as described previously [22]. Mice were tested prior to prostate antigen (PAg) injection (baseline) and after development of EAP. Referred tactile allodynia was evaluated using von Frey filaments applied to the lower abdomen [12]. Mice were placed individually in Plexiglas chambers (6 × 10 × 12 cm) on a stainless-steel wire grid floor and allowed to acclimate for 20 min. Testing was performed under standardized conditions (fixed time of day, single experimenter, blinded to group). Five filaments (0.04, 0.16, 0.4, 1.0, and 4.0 g; Stoelting, USA) were applied in ascending order of force. Each filament was applied for 1–2 s with 5 s inter-stimulus intervals, for a total of 10 applications per filament. Stimulation was confined to the lower abdomen overlying the prostate. A positive response was defined as one of the following: (1) sharp retraction of the abdomen, (2) immediate licking or scratching at the stimulus site, or (3) jumping. Response frequency was calculated as the percentage of positive responses out of 10 stimulations (e.g., 5/10 = 50%). Data are expressed as mean percentage response frequency ± SEM. For calculation of percentage change in responses from baseline (or pre-treatment baseline) the total responses of all five filaments were summed and the percent increase over baseline was calculated.

### Additional Behavioral tests

The animals were acclimated to the testing room environment for at least 2 days prior to the first test. The tests were performed during the light cycle. Animals had free access to food and water. Each test was video recorded by a digital camera connected to ANY-maze software (Stoelting Co, IL, USA). The data were analyzed using ANY-maze software. GraphPad Prism software (GraphPad Software, MA, USA) was used for statistical analysis. The tests were performed in this order: Dark and Light Box, Y maze, Novel Object Recognition and Open Field.

*Dark and Light Box test:* The animals are tested in a 40 x 40 cm box made of black plexiglass. This box is divided into 2 equal-sized chambers which are connected by a small opening, so the studied mouse can freely move between the 2 chambers. One of the chambers is covered (and dark) while the other is exposed to the light. The covered chamber is normally preferred by mice. A mouse is placed into the open side and the test lasts 5 minutes. We measured how much time the mouse spends in each chamber and the distance travelled in the light side. The preference for the light side is expressed as percentage: (time spent in light chamber/total testing time) x 100.

*Y-maze spontaneous alteration test:* Testing was done in a Y-shaped maze with three - high-walled (10 cm tall) arms at a 120° angle from each other. After being placed in the center of the maze, the mouse was allowed to freely explore the three arms for 5 minutes. Over the course of multiple arm entries, mice show a tendency to enter a less recently visited arm [13]. An entry is defined when all four limbs are within the arm. The sequence of arm entries is recorded, and the number of spontaneous alterations is calculated. A spontaneous alteration occurs when the mouse enters different arms in 3 consecutive entries. results are presented as percentage of correct spontaneous alterations: (number of the spontaneous alterations/the total number of entries – 2) × 100.

*Novel object recognition test (NOR):* NOR is a classic test of episodic memory [9]. Mice are habituated to an empty boxed arena (50 x 50 cm) for 1 hr. On the next day, during the acquisition trial, mice are allowed to explore two identical objects for 10 min and are then returned to the home cage. On the next day, for the NOR retention trial, mice are exposed to one familiar object and one novel object. We quantified the fraction of time spent exploring each object in each trial, and the distance traveled. The Discrimination index is expressed as: (time spent exploring the novel object – time spent exploring the familiar object)/total exploration time.

*Open Field test:* This test is widely used to assess exploratory behavior and general activity in rodents. We placed the mouse under study in a square boxed arena (40 x 40 cm) for 5 minutes. We quantified total distance travelled, and the time spent in the arena center vs. wall-hugging (thigmotaxis), which is a commonly used index of anxiety [24]. The ANY-maze software dissects the total space into 9 equal-sized squares, which therefore identifies the center square. The preference for the central area is expressed as a percentage: (time spent in middle/total test time) x 100

### DRG explant culture with inflammatory stimulation

Ex vivo DRG explant cultures were prepared with minor modifications of published protocols [14; 23; 25; 26]. Briefly, 5–7-week-old C57BL/6 mice were euthanized with a lethal dose of isoflurane. Under aseptic conditions, the vertebral column from sacral to lumbar levels was exposed, and the spinal canal was opened along the midline. DRG (L2–S1) were dissected using #7 forceps, rinsed twice in sterile HBSS, and transferred to 96-well plates containing 200 µl DMEM supplemented with 10% FBS, 2% B27, and 10 ng/ml NGF. DRG were cultured at 37°C, 5% CO_2_ for 24 h with or without 100 ng/ml SELTA. To model in vitro inflammatory stimulation, DRG were treated with recombinant TNF-α (10 ng/ml) for 12-18 hours prior to experimental assays. This exposure paradigm was selected based on prior studies demonstrating a significant role for TNF-α signaling in dorsal root ganglion neurons following peripheral nerve injury [6; 10; 29] and the use of prolonged TNF-α treatment to model neuropathic and inflammatory pain–associated neuronal states in vitro [28; 31]. Chronic TNF-α exposure has been shown to induce transcriptional and protein-level changes in sensory neurons, including increased expression of the voltage-gated sodium channel NaV1.7, which is upregulated in models of nerve injury–associated pain [28].

### Primary DRG sensory neuron culture and *in vitro* inflammatory stimulation

DRG were dissected from C57BL/6 mice into cold HBSS and minced into ∼0.2-mm fragments. After centrifugation at 100 g for 1 min, tissue was digested with papain (20 min, 37°C) followed by collagenase/dispase (20 min, 37°C). Cells were centrifuged at 400 g for 4 min and resuspended in L-15 medium containing 5% FCS, penicillin/streptomycin, and HEPES. Neurons were enriched using a Percoll gradient (1300 g, 10 min), then resuspended in F12/DMEM containing 10% FCS, 10 ng/ml 2.5S NGF, penicillin/streptomycin, HEPES and 10μM Cytosine beta-D-arabinofuranoside (Ara-C). Cells were plated at 5 × 10³ neurons per well in 4-chamber glass-bottom dishes (D35C4-20-0-N; Cellvis) for subsequent experiments.

For modeling in vitro inflammatory stimulation, dissociated neurons were treated as above with recombinant TNF-α (10 ng/ml) for 12-18 hours prior to experimental assays. This exposure paradigm was selected based on prior studies demonstrating a significant role for TNF-α signaling in dorsal root ganglion neurons following peripheral nerve injury [6; 10; 29].

### Calcium imaging

Calcium imaging was performed as described previously [20]. DRG neuron cultures exposed to inflammatory stimulation were loaded with 1 µM Fura-2 AM and 1 µM Pluronic F-127 (Life Technologies) in HBSS (pH 7.4; 137 mM NaCl, 5.4 mM KCl, 1.3 mM CaCl_2_, 0.5 mM MgCl_2_, 0.4 mM MgSO_4_, 4.2 mM NaHCO_3_, 0.44 mM KH_2_PO_4_, 0.34 mM Na_2_HPO_4_, 5.5 mM glucose, 25 mM HEPES, 1 mg/ml BSA) at 37°C for 30 min, then washed in HBSS. For SELTA experiments, neurons were incubated with 100ng/ml of SELTA alone in HBSS, SELTA plus rat anti-PD-1 neutralizing antibody, or SELTA plus PD-1 control peptide and anti-PD-1 antibody and returned to the incubator for ≥10 min before recording. All groups were treated for the same period. Cells were transferred to the imaging setup and equilibrated to room temperature. Baseline intracellular Ca^2+^ levels were recorded for at least 60 s using ratiometric imaging at 340/380-nm excitation, followed by ATP stimulation. Signals were recorded for up to 600 s. At the end of each experiment, 2 µM ionomycin was applied to elicit a maximal Ca^2+^ response and confirm cell viability. For each chamber, ∼10–80 cells were imaged using a Leica DMI6000B microscope with a 40× oil-immersion objective and experiments were repeated three times with independent cultures. Images were captured with a Hamamatsu CCD camera and analyzed using MetaFluor (Molecular Devices). F340/F380 ratios were used to quantify Ca^2+^ responses.

### RT-PCR and qPCR

RNA was isolated from iliac lymph nodes, splenocytes, and cultured sensory neurons as described [17]. RNA concentration was measured using a Spark microplate reader (Tecan). cDNA was synthesized with SuperScript® III First-Strand Synthesis System (Invitrogen, Cat. 18064014) according to the manufacturer’s protocol. qRT-PCR was performed using SsoAdvanced^TM^ Universal SYBR® Green Supermix (Bio-Rad, Cat. 1725271) with 50 ng cDNA per 25-µl reaction on a CFX Connect™ Real-Time PCR Detection System (Bio-Rad). Primers were designed using the IDT PrimerQuest™ Tool. Relative expression was calculated using the ΔΔCt method with appropriate internal controls. Samples were run in biological triplicate (3–4 mice per group) with technical replicates. Primer sequences for full-length Pdcd1 are listed in Table 1.

**Table 1:**
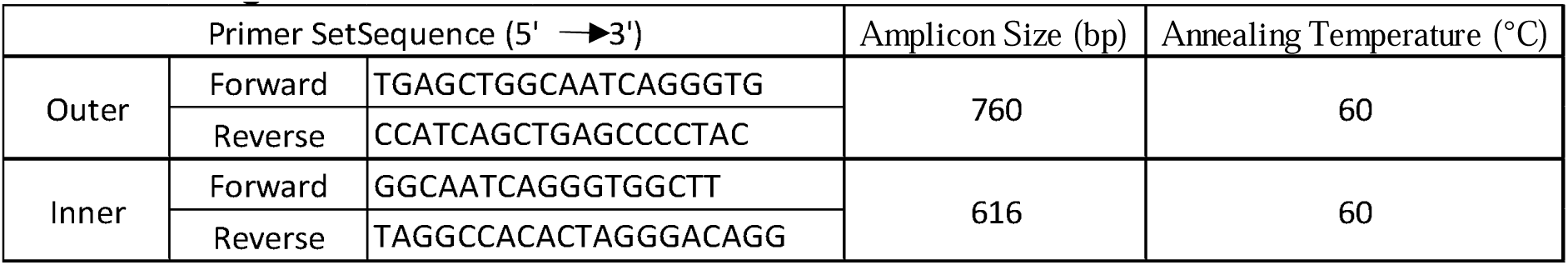
Primer Sequences and PCR Conditions for Nested PCR Amplification of the Full-Length PD-1 Gene.

### Immunohistochemistry and immunofluorescence

Mice were anesthetized with 2.5% isoflurane (Isothesia; Butler) and transcardially perfused with cold PBS followed by 4% paraformaldehyde (PFA). DRG were removed and post-fixed in 4% PFA (DRG, 1 h), then cryoprotected in 30% sucrose overnight. Tissues were frozen in dry ice–chilled isopentane, embedded in OCT (Tissue-Tek), and stored at −80°C. L2–S1 DRG and prostate were sectioned at 10 µm on a cryostat (Leica), mounted on Superfrost Plus slides, and stored at −80°C. Sections were fixed in 4% PFA for 10 min, subjected to antigen retrieval in 10 mM sodium citrate buffer (pH 9.0) for 10 min at 80°C, and blocked in 0.3% Triton X-100/5% donkey serum in PBS for 30 min. Primary antibodies were applied overnight at 4°C in blocking buffer, followed by three 10-min PBS washes and 2-h incubation with Alexa Fluor 405-, 488-, 555-, or 647-conjugated secondary antibodies. Nuclear counterstaining was performed with Hoechst 33342 or DAPI, and sections were mounted in ProLong Diamond Antifade (Invitrogen). Primary antibodies included rabbit anti-PD-1 (Sigma-Aldrich, Cat. PRS4065), rabbit anti-phospho-PD-1 (Abcam, Cat. ab206378), mouse anti-CGRP (Abcam, Cat. ab81887), rabbit anti-CGRP (Cell Signaling, Cat. 14959S), goat anti-IBA1 (Novus, Cat. NB100-10288SS), guinea pig anti-NeuN (Synaptic Systems, Cat. 266004), chicken anti-βIII-tubulin (Novus, Cat. NB100-1612), goat anti-TLR2 (Santa Cruz, Cat. SC-12504), mouse anti-TLR2 (Novus, Cat. NBP2-30097), and mouse anti-lipoteichoic acid (Thermo Fisher, Cat. MA1-40134).

### Confocal imaging and quantitative immunofluorescence

Images were acquired on a Nikon A1 confocal microscope using Plan Apo 20× (NA 0.75) and 60× oil (NA 1.4) objectives. For quantitative analysis, comparable tissue regions were selected, and all groups were imaged using identical confocal settings established from a reference template. After setting pixel resolution and line averaging, laser power and detector gain for each channel were optimized and then kept constant across samples. Following noise reduction and thresholding of the NeuN-immunoreactive channel in ImageJ, regions of interest (ROIs) corresponding to neuronal somata were defined with manual correction of cell borders. The same ROIs were applied to PD-1 and phospho-PD-1 channels to extract mean fluorescence intensity. For cell size–dependent analyses, DRG neurons were classified as small (<300 µm^2^), medium (300–500 µm^2^), or large (>500 µm^2^).

### Statistical analysis

Statistical analyses were performed using GraphPad Prism^TM^ and Microsoft Excel^TM^. Data are presented as mean ± SEM unless otherwise indicated. Comparisons between two groups used Student’s t test. Comparisons involving more than two groups used one-way or two-way ANOVA, as appropriate, followed by Tukey’s or Bonferroni’s post hoc tests. A p value < 0.05 was considered statistically significant.

### Use of artificial intelligence tools

Portions of the manuscript text were edited for clarity and language using a large language model (ChatGPT, OpenAI). The tool was used exclusively for language refinement and formatting. All scientific content, data analysis, interpretation, and conclusions were developed and verified by the authors, who take full responsibility for the integrity and accuracy of the work.

## Results

### SELTA activates TLR2/6-dependent NF-κB signaling and associates with sensory neurons

To identify receptor usage by SELTA, we examined NF-κB activity in HEK293 reporter cells expressing human TLR2 heterodimers. Cells expressing hTLR2 together with both hTLR1 and hTLR6, hTLR2/1, hTLR2/6, or vector control (HEK-Null) were stimulated with Pam2CSK4, Pam3CSK4, or SELTA (10 or 100 ng/mL) for 24 hours. As expected, Pam2CSK4 preferentially activated TLR2/6-expressing cells and Pam3CSK4 preferentially activated TLR2/1-expressing cells, confirming functional receptor expression (Fig. 1A; Supplementary Fig. S1). SELTA induced robust NF-κB activation in HEK-hTLR2/6 cells and in cells expressing hTLR2 with both hTLR1 and hTLR6, with minimal activation in HEK-hTLR2/1 or HEK-Null controls (Fig. 1A). Two-way ANOVA with Tukey’s multiple comparisons demonstrated significantly greater activation in HEK-hTLR2/6 compared with HEK-hTLR2/1 within each SELTA condition. Analysis of the dose–response of SELTA (0.001–1000 ng/mL) showed a concentration-dependent increase in NF-κB activity in HEK-hTLR2/6 and HEK-hTLR2 cells, whereas HEK-hTLR2/1 and HEK-Null cells showed negligible activation across concentrations (Fig. 1B). Significant increases were observed in HEK-hTLR2/6 cells at 100 and 1000 ng/mL compared with 10 ng/mL. These findings indicate that SELTA activates NF-κB signaling in a TLR2/6-dependent context.

**Figure 1.**
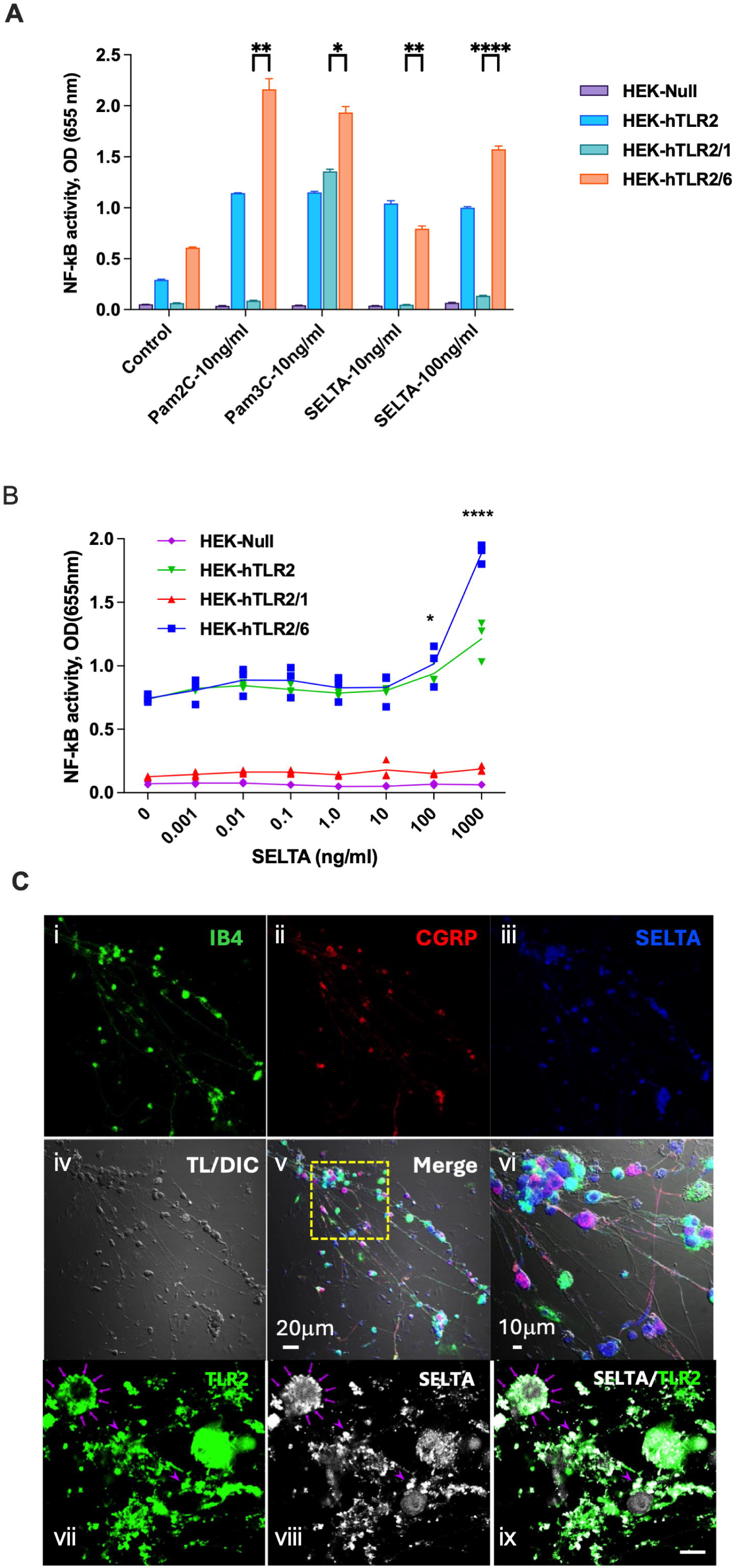
SELTA selectively activates TLR2/6 signaling and engages sensory neurons. (A) HEK293 NF-κB reporter cells expressing hTLR2 with both hTLR1 and hTLR6, hTLR2/1, hTLR2/6, or vector control (HEK-Null) were stimulated with Pam2CSK4 (TLR2/6 agonist), Pam3CSK4 (TLR2/1 agonist), or 10ng/ml and 100ng/ml SELTA for 24 hours. NF-κB activation was quantified by SEAP reporter activity (OD 655 nm). SELTA induced robust activation in HEK-hTLR2/6 and hTLR2 cells with minimal activity in HEK-hTLR2/1 cells. Data are mean ± SEM from three independent experiments. Statistical analysis was performed using two-way ANOVA with Tukey’s multiple comparisons test. Asterisks denote significance of HEK-hTLR2/6 relative to HEK-hTLR2/1 within each stimulus condition (*p<0.05, **p<0.01, ***p<0.001, ****p<0.0001). (B) Dose–response analysis of SELTA (0.001–1000 ng/mL) in HEK reporter lines. SELTA produced a concentration-dependent increase in NF-κB activity selectively in HEK-hTLR2/6 and HEK-hTLR2, with negligible activation of HEK-hTLR2/1 or HEK-Null cells. Data are mean ± SEM from three experiments; significance determined by two-way ANOVA with multiple comparisons (*p<0.05, ****p<0.0001 at 100ng/ml and 1000ng/ml vs 10ng/ml). (C) Confocal Immunofluorescence imaging of mouse primary dorsal root ganglia neurons treated with SELTA. Neurons were stained for IB4 (i, green), CGRP (ii, red), and SELTA (iii, blue). Differential interference contrast (DIC) imaging (iv) demonstrates neuronal morphology. Merged images (v–vi) show SELTA association with both IB4-positive non-peptidergic and CGRP-positive peptidergic neurons. Insets depict higher magnification views (scale bars: 20 μm and 10 μm). (vii–ix) Confocal imaging of TLR2 (green), SELTA (white), and merged channels in mouse primary dorsal root ganglia neurons. SELTA puncta (purple arrows) colocalize with TLR2-positive structures, consistent with receptor engagement at the neuronal surface. Representative images from ≥3 independent preparations are shown. Scale bar is 10 μm.

We next examined SELTA localization in dissociated mouse primary dorsal root ganglia (DRG) neurons. Confocal imaging of neurons demonstrated SELTA immunoreactivity in IB4-positive and CGRP-positive neuronal subsets (Fig. 1Ci-iii). Merged images with DIC (iv-vi) revealed SELTA labeling along neuronal somata and processes.

We next examined whether the TLR2 receptor is expressed on cultured primary neurons and if SELTA was co-localized with TLR2. We first validated the specificity of TLR2 antibody staining using confocal microscopy in primary DRG neurons with and without anti-TLR2 primary antibody (Supplementary Figs. S2). Co-staining was subsequently performed using this antibody along with an anti-LTA antibody on primary DRG cultures that had been exposed to 100ng/ml SELTA for 24 hours prior to analysis. Co-labeling observed as overlapping signal was detected between SELTA and TLR2 (Fig. 1C vii-ix) suggesting similar sites of localization.

Together, these data show that SELTA activates NF-κB signaling in TLR2/6-expressing reporter cells and is detected in TLR2-positive sensory neurons in dorsal root ganglia.

### SELTA increases neuronal PD-1 expression in sensory ganglia

To examine the distribution of PD-1 across different tissues in mice, we first assessed *Pdcd1* transcript levels by RT-PCR. Baseline *Pdcd1* expression was detected in immune organs (spleen, lymph nodes) and prostate, as well as throughout the sensory and central nervous system, including dorsal root ganglia (DRG), spinal cord, thalamus, hippocampus, and cortex (Fig. 2A). To determine whether SELTA can modulate PD-1 levels, we evaluated PD-1 mRNA expression in an ex vivo DRG preparation. Isolated DRG were treated with recombinant TNF-α (10 ng/ml) for 12-18 hours prior to experimental assays. This exposure paradigm was selected based on prior studies demonstrating the use of prolonged TNF-α treatment to model neuropathic and inflammatory pain–associated neuronal states in vitro [28; 31] and a significant role for TNF-α signaling in dorsal root ganglion neurons following peripheral nerve injury [6; 10; 29]. Using this *in vitro* inflammatory model, lumbosacral DRG isolated from mice treated with 100ng/ml SELTA for 24 hours and RNA was isolated. QPCR analysis was performed and demonstrated that *Pdcd1* mRNA relative to GAPDH was significantly increased in SELTA-treated DRGs compared with controls (Fig. 2B).

**Figure 2.**
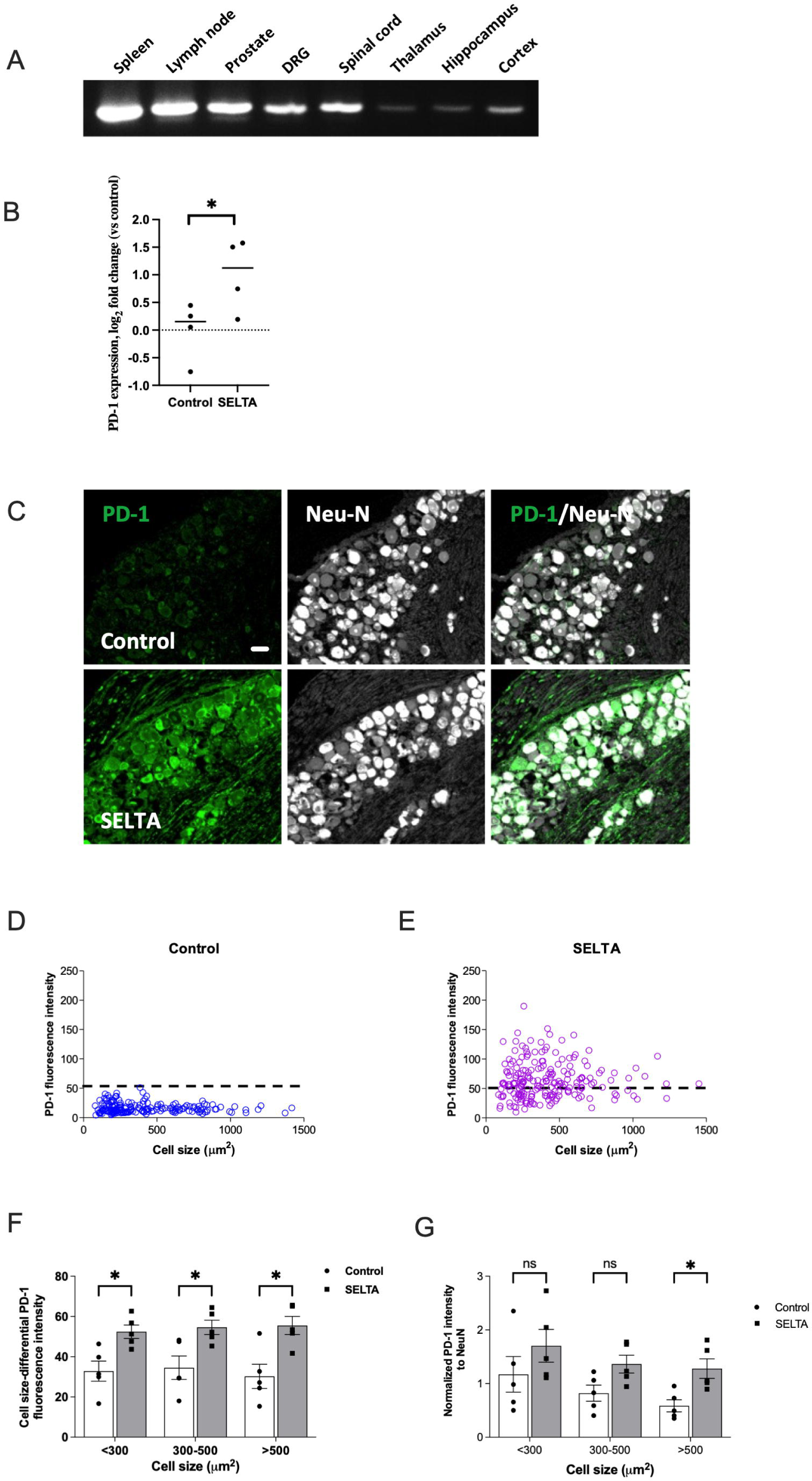
Basal neuronal PD-1 expression and SELTA-induced changes. (A) RT-PCR analysis of *Pdcd1* (PD-1) mRNA expression across splenocytes, lymph nodes, prostate, dorsal root ganglia (DRG), spinal cord, thalamus, hippocampus, and cortex. (B) Quantitative PCR analysis of *Pdcd1* mRNA expression in DRG treated with SELTA (100 ng/mL, 24 h) or vehicle control in vitro. Expression was normalized to GAPDH and calculated using the 2^−ΔΔCt method. Data are presented as log2 fold change relative to control. Statistical analysis was performed on ΔCt values; *p < 0.05. (C) Ex vivo DRG cultures showing PD-1 immunoreactivity in untreated (control) and SELTA-treated (100 ng/mL, 24 h) neurons. PD-1 (green) and NeuN (white) staining identify neuronal PD-1 expression. Scale bar = 30 µm. (D&E) Distribution of PD-1 immunoreactive intensity across neuronal size classes in representative confocal images of control and SELTA-treated DRG. (F) Quantification of overall PD-1 immunoreactivity in DRG neurons from control and SELTA-treated mice. (G) Quantification of PD-1 immunoreactivity normalized to NeuN in DRG neurons from control and SELTA-treated mice. For all quantitative analyses, data were collected from 1321 and 1251 individual neurons in control and SELTA-treated DRG, respectively (n=5 mice per group). Student’s *t*-test was used for two-group comparisons and two-way ANOVA for analyses involving multiple groups. \**P* < 0.05; Abbreviations: NeuN, neuronal nuclear antigen; DRG, dorsal root ganglion; ANOVA, analysis of variance.

To determine whether the modulation of PD-1 mRNA expression extends to protein levels, we evaluated PD-1 immunoreactivity in the *ex vivo* DRG preparation. We first validated an anti-PD-1 antibody for immunostaining specificity to neuronal PD-1 by demonstrating binding to NeuN expressing neurons using immunofluorescence with or without primary antibody present (Supplementary Fig. S2). Subsequent studies using SELTA treatment (100 ng/mL, 24 h) in this model showed increased PD-1 signal intensity relative to untreated cultures (Fig. 2C). Stratification by neuronal size showed that SELTA elevated PD-1 across small, medium, and large-diameter neurons (Fig. 2D&E). Quantification of PD-1 immunoreactivity in individual neurons confirmed a significant increase in overall intensity in all three neuronal subtypes (Fig. 2F).

Normalization to individual NeuN immunostaining intensity showed that significance was limited to large-diameter neurons (Fig. 2G). Taken together, these results show that SELTA enhances PD-1 expression in sensory ganglia under inflammatory conditions, at both transcript and protein levels.

### SELTA exposure increases PD-1 phosphorylation and reduces ATP-evoked calcium signaling in DRG neurons

Since SELTA increased total PD-1 expression in DRG neurons, we next examined whether SELTA can elicit PD-1 activation. Utilizing an antibody that binds phosphorylated tyrosine 248 (Y248) in the C-terminal tail of PD-1 which is associated with activation of PD-1 inhibitory signaling, we sought to understand the effect of SELTA. We first validated the anti-PD-1 antibody specificity using DRG staining with and without the primary antibody in conjunction with NeuN (Supplementary Fig. S2). Using this *ex vivo* DRG preparation, SELTA treatment (100 ng/mL, 24 h) significantly increased pPD-1 immunoreactivity compared with untreated DRG (Fig. 3A). This effect was observed to be statistically significant when measured across all neurons (Fig. 3B) and showed an increase that was not statistically significant when normalized against NeuN (Fig. 3C). These findings suggest that SELTA can elicit PD-1 activation in DRG neurons.

**Figure 3.**
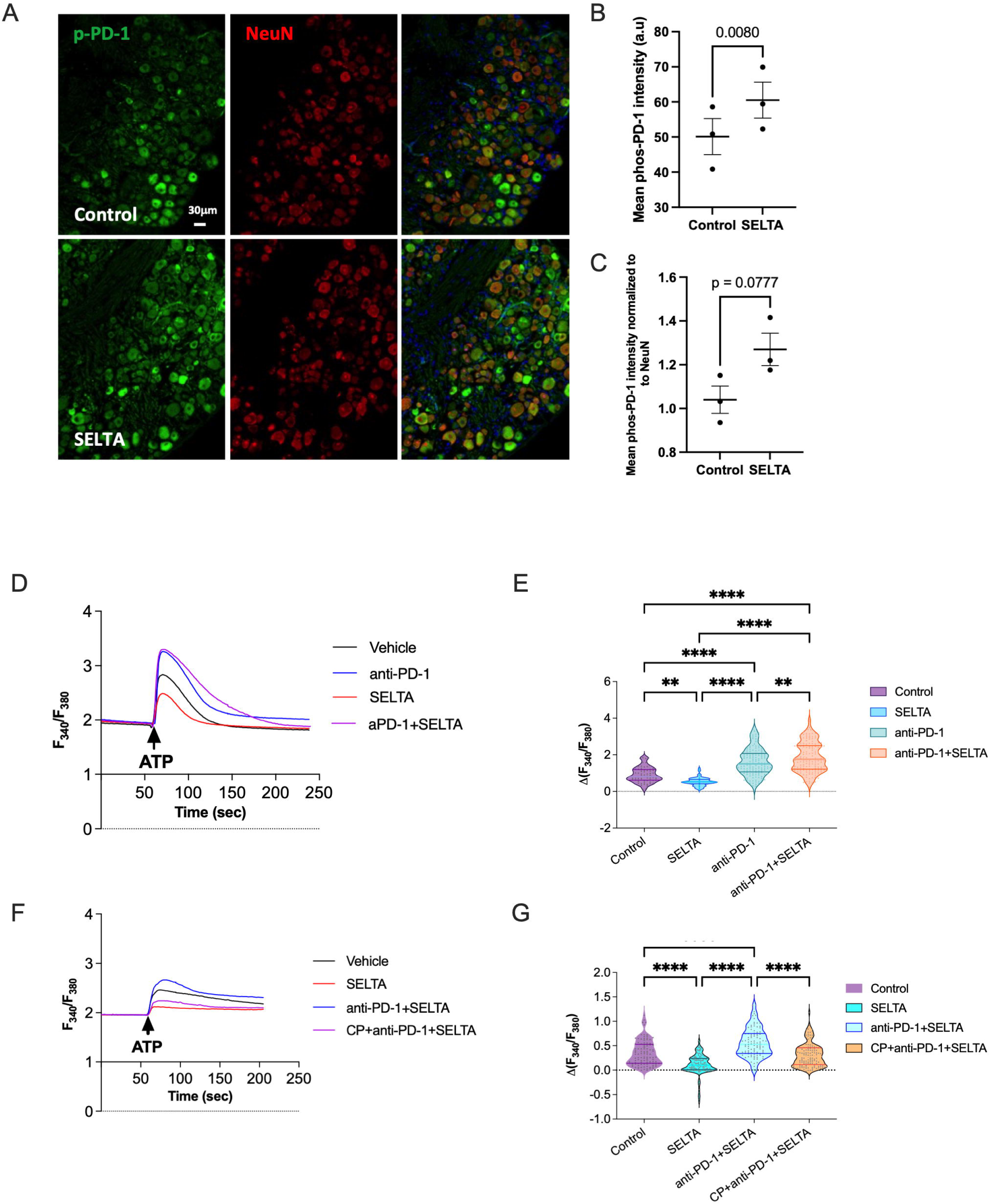
SELTA enhances PD-1 phosphorylation at Y248 and inhibits ATP-evoked Ca^2+^responses in a PD-1 dependent manner. (A) Representative confocal images in *ex vivo* DRG cultures showing pPD-1 (green) and NeuN (red) immunoreactivity in control (top) and SELTA-treated (100 ng/mL, 24 h; bottom) preparations. Scale bar = 30 µm. (B) Quantification of pPD-1 immunoreactivity across all neurons and (C) pPD-1 immunoreactivity normalized to NeuN. A total of 585 control neurons and 807 SELTA-treated neurons were analyzed from 3 paired mice. For statistical analysis, neuronal measurements were averaged within each mouse and condition, and paired comparisons were performed using animal-level means. Student’s *t*-test was used for two-group comparisons, and *P* values are shown. (D–G) For intracellular calcium studies primary DRG sensory neurons were grown on coverslips, loaded with Fura-2 and stimulated with 20 µM ATP followed by measurement of change in intracellular calcium levels using the F340/F380 fluorescence ratio. Baseline values were recorded for at least 60 seconds before ATP application. Peak ATP-evoked calcium responses were calculated as the difference between baseline and the average maximal F340/F380 ratio within the 10-second post-stimulation window. SELTA pretreatment (15-30 minutes) significantly reduced ATP-induced intracellular Ca^2+^ elevation compared with untreated controls (D, E). This inhibitory effect was abolished when neurons were pretreated with a PD-1 neutralizing antibody (anti-PD-1, 30 minutes), (D, E). When the PD-1 neutralizing antibody was pre-incubated with a PD-1 control peptide (CP + anti-PD-1 + SELTA), antibody neutralization was blocked, and SELTA’s inhibitory effect on ATP-evoked Ca^2+^ responses was restored (F, G). Ca^2+^ imaging data were obtained from three independent experiments, with a total of 67–147 neurons analyzed per group. Primary neuronal cultures were prepared from lumbar DRG pooled from 6–8 mice aged 10–15 days. Student’s *t*-test was used for comparisons between two groups, and one-way ANOVA with multiple comparisons were used for analyses involving more than two groups. **P < 0.01, and ****P < 0.0001. Abbreviations: pPD-1, phosphorylated programmed cell death-1; NeuN, neuronal nuclear antigen

To determine whether SELTA-induced signaling can affect activation of DRG neurons, we studied ATP-induced intracellular calcium responses using real-time ratiometric imaging with Fura-2. SELTA pretreatment significantly reduced ATP-induced intracellular Ca^2+^ elevations compared with untreated controls (Fig. 3D, E). This inhibitory effect was abolished when neurons were pretreated with a PD-1 neutralizing antibody (anti-PD-1), demonstrating that SELTA’s calcium-suppressive effect requires intact PD-1 signaling (Fig. 3D, E). When the PD-1 neutralizing antibody was pre-incubated with a PD-1 control (immunizing) peptide (CP + anti-PD-1 + SELTA), antibody neutralization was blocked, and SELTA’s inhibitory effect on ATP-evoked Ca^2+^ responses was restored (Fig. 3F, G). These results show that SELTA functionally suppresses neuronal calcium mobilization, a key marker of sensory neuron activation, in a PD-1–dependent manner.

### SELTA attenuates pelvic hypersensitivity in EAP mice in a concentration-dependent manner

We next examined the effect of SELTA in experimental autoimmune prostatitis (EAP), an *in vivo* model of chronic pelvic pain. Induction of EAP results in a progressive increase in allodynia/hyperalgesia from days 0-28 in the suprapubic region in mice that subsequently stays elevated (Fig. 4A-C). Mice received intraurethral administration of 100 ng SELTA or saline one day prior to von Frey testing. Therapeutic efficacy was assessed on days 28, 35, and 42, and then weekly thereafter. A second intraurethral dose of 100 ng SELTA was administered on day 70, followed by continued behavioral testing through day 119 (Fig 4A and Supplementary Fig. S3). The first SELTA administration significantly reduced pelvic allodynia compared with untreated EAP controls, with improvements evident by day 28 and staying significantly reduced through day 42 (Fig. 4B). This analgesic effect gradually diminished, peaking by days 70 at which time a second SELTA instillation was performed resulting in a significant second reduction in pelvic hypersensitivity (Fig. 4C).

**Figure 4.**
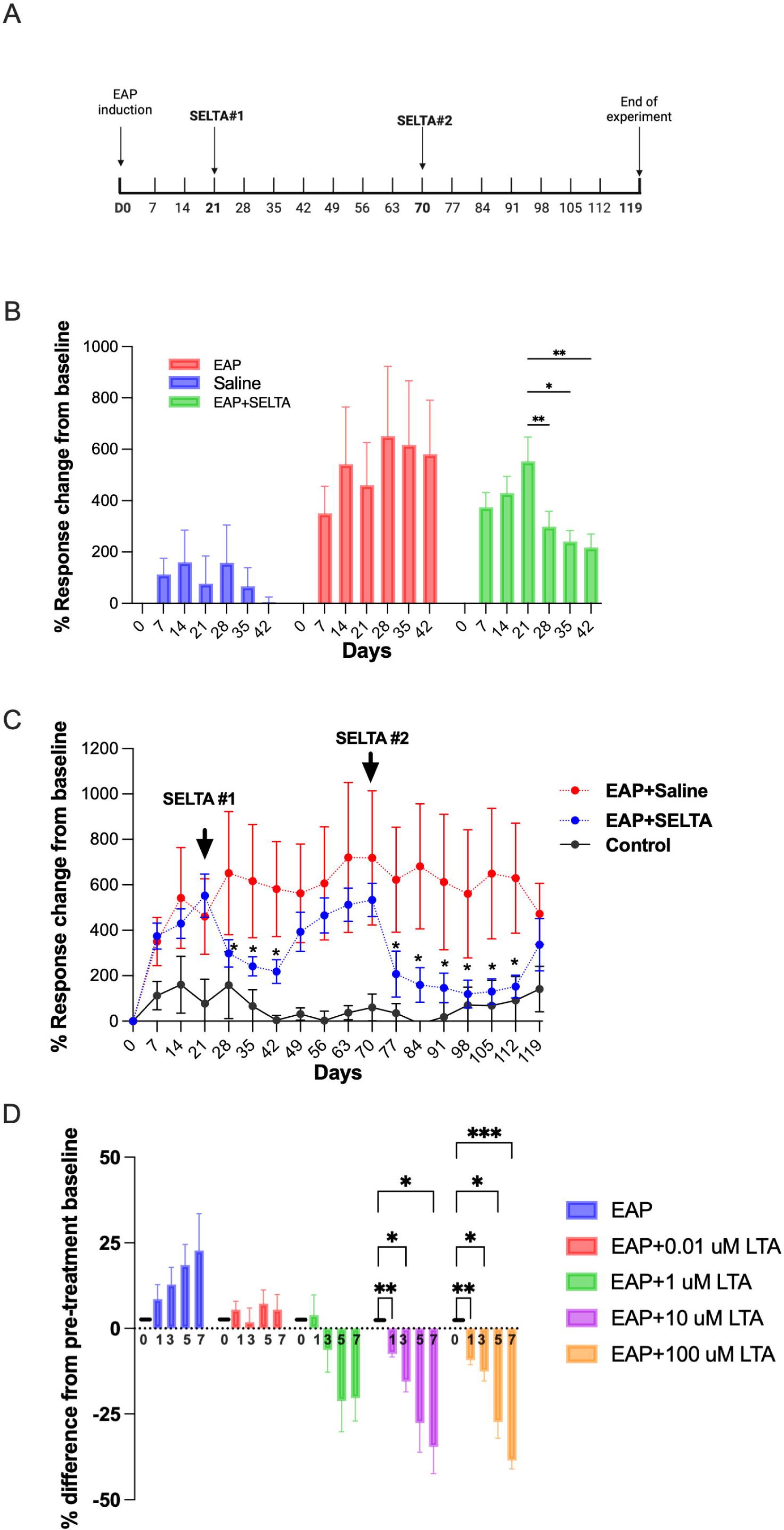
SELTA reproducibly attenuates pelvic allodynia in EAP mice with both and shows a concentration-dependent effect. (A) Schematic of the experimental design illustrating induction of experimental autoimmune prostatitis (EAP), timing of weekly behavioral assessments, and SELTA administration. SELTA or saline (100ng in 10 μl) was delivered intraurethrally on days 21 and 70. (B) Early-phase behavioral response extracted from panel C, showing pelvic tactile allodynia over the first 42 days and highlighting the effect of the first SELTA instillation at day 21. (C) Long-term pelvic tactile allodynia in saline-treated mice, EAP mice, and EAP mice receiving SELTA on days 21 and 70. Behavioral responses were assessed every 7 days up to day 119 and expressed as percentage change in response frequency relative to baseline. (D) Single-dose intraurethral SELTA dose–response study (0.01, 1, 10, or 100 μM corresponding to 0.81 ng, 81 ng, 810 ng, and 8.1 µg, respectively) in EAP mice. Pelvic tactile allodynia was monitored over 7 days following treatment, and the results are displayed as percent difference relative to untreated EAP controls. For all behavioral analyses, two-way ANOVA with Tukey’s multiple comparisons test was used. *P* < 0.05; P < 0.01; *P* < 0.001. Group sizes were 5–10 mice per condition. Abbreviations: EAP, experimental autoimmune prostatitis; SELTA, staphylococcal lipoteichoic acid; ANOVA, analysis of variance.

To assess concentration dependence, EAP mice received a single intraurethral SELTA dose a day prior to day 21 across a 10,000-fold concentration range. While EAP mice showed an expected increase in pelvic hypersensitivity over the next 7 days, SELTA-treated mice showed a graded reduction in pelvic hypersensitivity, with higher concentrations of SELTA (10 and 100 μM) producing significantly greater attenuation (Fig. 4D and Supplementary Fig. S4). These findings show that SELTA reduces EAP-induced pelvic allodynia after both single and repeat dosing and that its analgesic activity is concentration dependent.

### Sensory neuron–specific deletion of PD-1 reduces SELTA-induced analgesia in EAP mice

To determine whether neuronal PD-1 is required for SELTA-mediated analgesia, we generated sensory neuron–specific PD-1 conditional knockout (CKO) mice by crossing *Pdcd1-*floxed mice with Advillin^Cre^ drivers (Supplementary Fig. S5A). Confocal imaging of DRG sections from PD-1^f/f^ and PD-1^fl/fl^;Advillin^Cre^ mice were used to validate the anti-PD-1 antibody using pre-incubation with a control peptide (Supplementary Fig. S5B) and demonstrated a reduction of PD-I immunoreactivity specifically in NeuN-positive DRG neurons of PD-1^fl/fl^;Advillin^Cre^ mice. In contrast, immunoreactivity for PD-1 expression was retained in IBA-1–positive macrophages/glia in the same sections (Supplementary Fig. S5B). To study baseline functional impact of PD-1 deletion, behavioral assessments were performed using dark and light box (Supplementary Fig. S7A), Y Maze (Supplementary Fig. S7B), novel object recognition (Supplementary Fig. S7C) and open field (Supplementary Fig. S7D) assays. No significant differences were observed between PD-1^fl/fl^ and PD-1^fl/fl^;Advillin^Cre^ mice in any of these assays ruling out any impact of neuronal PD-1 reduction on anxiety-like behavior, locomotor activity, exploratory behavior, or recognition memory.

EAP was induced in both genotypes and allowed to develop for 50 days before intraurethral SELTA administration (100ng, Fig. 5A). In PD-1 mice, SELTA produced reductions in pelvic tactile allodynia at 1-, 3-, 5-, and 7-days post-treatment (Fig. 5A, Band Supplementary Fig. S8A). In contrast, PD-1^fl/fl^;Advillin^Cre^ mice showed no significant reduction from pretreatment baseline at days 1, 3, 5 and 7 following SELTA instillation (Fig. 5B and Supplementary Fig. S8B). Together, these data suggest a requirement for PD-1 expression in advillin-expressing neurons for SELTA-induced analgesia.

**Figure 5.**
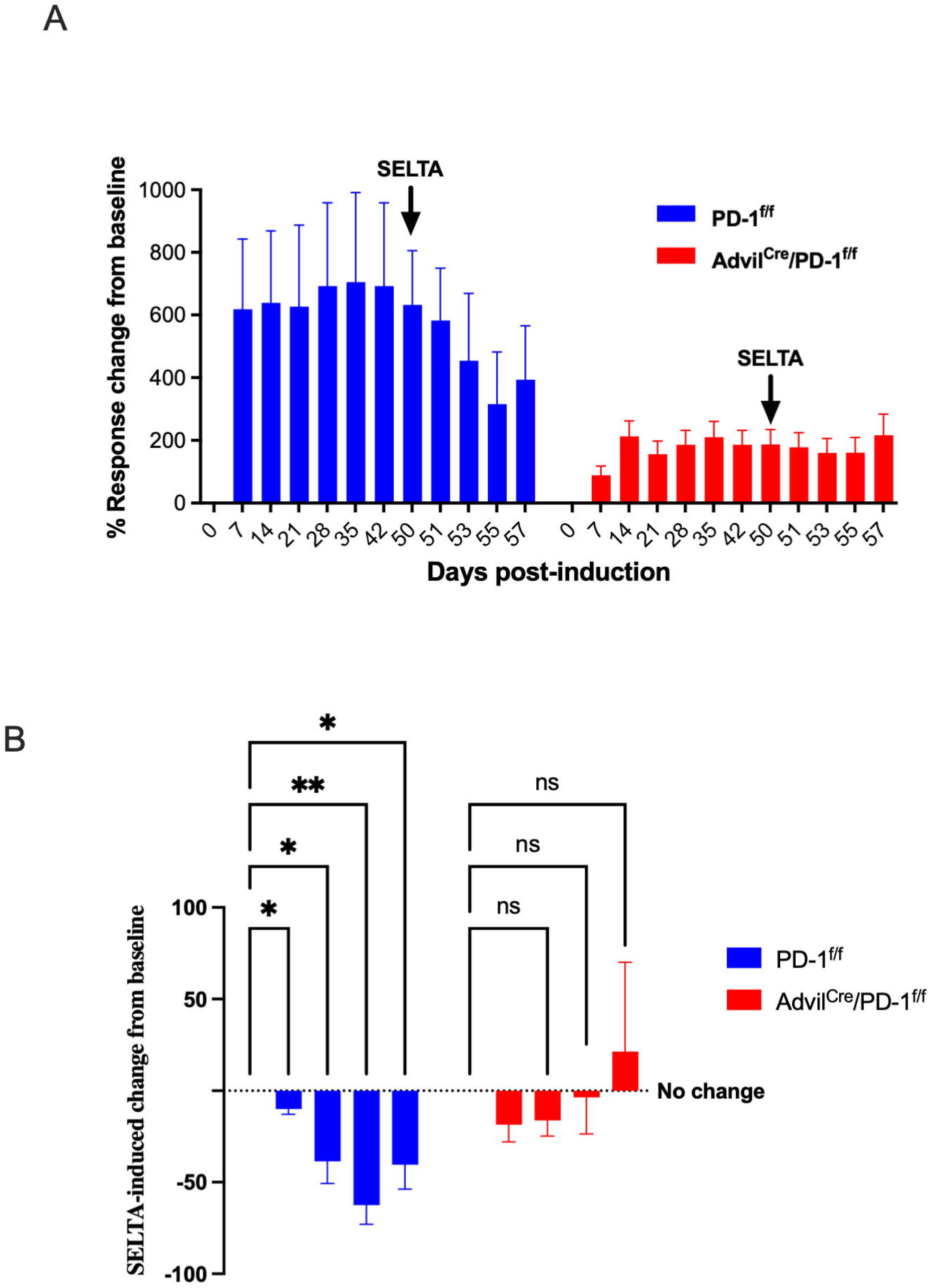
Neuronal PD-1 is required for SELTA-induced attenuation of pelvic allodynia in EAP mice. (A) Pelvic tactile allodynia in EAP-treated PD-1^fl/fl^ and PD-1^fl/fl^;Advillin^Cre^ mice. EAP was allowed to develop for 50 days before mice received a single intraurethral SELTA instillation (100ng). Pelvic tactile hypersensitivity was then assessed at 1, 3, 5, and 7 days after SELTA treatment using a graded series of von Frey filaments. Data are expressed as percentage increase in total response frequency relative to baseline. Arrows indicate the day of SELTA instillation after collection of a pre-treatment baseline. (B) Comparison of SELTA efficacy between PD-1^fl/fl^ and PD-1^fl/fl^;Advillin^Cre^ genotypes, displayed as the percent difference in SELTA-induced change from pre-treatment baseline across the 1–7-day (equal to the window from day-50 to day-57) post-treatment window. For statistical analysis, two-way ANOVA was used for analyses involving more than two groups. \**P* < 0.05; \*\**P* < 0.01. *n* = 9-11 mice per group. Abbreviations: EAP, experimental autoimmune prostatitis; SELTA, staphylococcal lipoteichoic acid analog; ANOVA, analysis of variance.

### SELTA-induced analgesia requires PD-1 expression in both sensory neurons and CD4^+^T cells

While deletion of PD-1 in sensory neurons attenuated SELTA efficacy (Fig. 5A), this finding does not exclude contributions from other PD-1–expressing cell populations. To evaluate the role of immune cell PD-1, we generated mice with conditional deletion of PD-1 in CD4^+^ T cells (CD4^Cre^; PD-1^fl/fl^) and mice with combined deletion in advillin-expressing sensory neurons and CD4⁺ T cells (Advillin^Cre^;CD4^Cre^ PD-1^fl/fl^).

Successful recombination in each line was confirmed by PCR genotyping (Supplementary Fig. S6A). Loss of PD-1 expression in CD4⁺ T cells was validated by immunofluorescence in lymph node sections demonstrating reduced PD-1 immunoreactivity within CD4⁺ regions, and by flow cytometric analysis of splenocytes showing decreased PD-1 expression in CD4⁺ T cells (Supplementary Fig. S6B–C). Generation of the sensory neuron–specific PD-1 conditional knockout was confirmed using the Advillin^Cre^ system (Supplementary Fig. S6A). PD-1 antibody specificity was verified using peptide blocking controls, and selective reduction of PD-1 immunoreactivity in NeuN-positive dorsal root ganglion (DRG) neurons, but not IBA-1–positive cells confirmed neuron-specific knockdown (Supplementary Fig. S5B).

Experimental autoimmune prostatitis (EAP) was induced in CD4^Cre^; PD-1^fl/fl^, Advillin^Cre^; PD-1^fl/fl^, Advillin^Cre^;CD4^Cre^; PD-1^fl/fl^, and wild-type littermate controls. Across all genotypes, EAP induction was associated with progressive reductions in nociceptive thresholds from days 0–21 prior to SELTA treatment, indicating comparable development of pelvic hypersensitivity (Fig. 6A and Supplementary Fig. S9). Following intraurethral administration of 100 ng SELTA, wild-type mice exhibited a significant reduction in pelvic hypersensitivity relative to pretreatment baseline. In contrast, mice lacking PD-1 in advillin-expressing sensory neurons or in CD4^+^ T cells showed markedly attenuated responses, with persistent hypersensitivity through day 28 (Fig. 6A). Deletion of PD-1 in both neuronal and immune compartments did not further reduce the response beyond that observed in either single conditional knockout.

**Figure 6.**
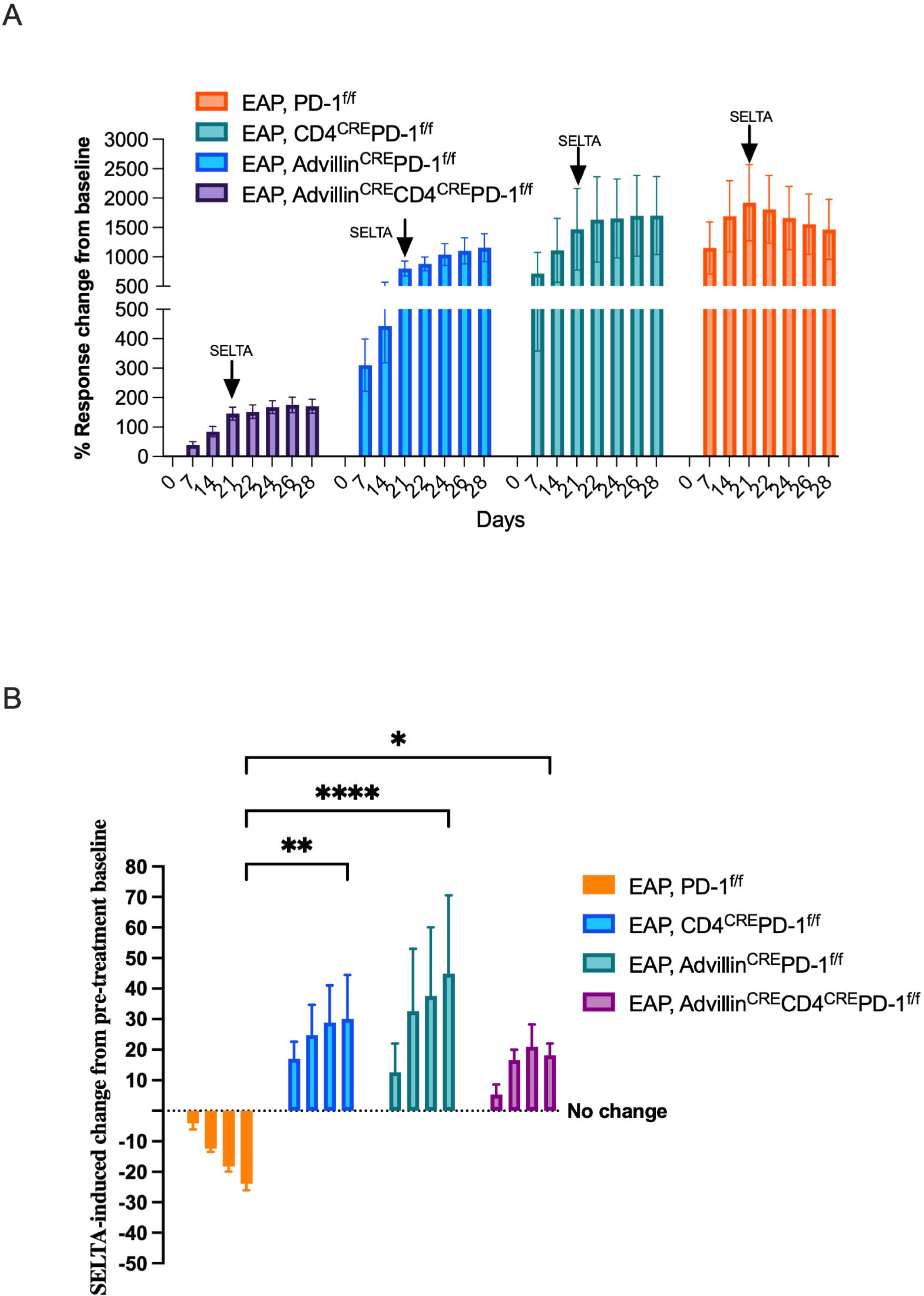
SELTA-induced analgesia requires PD-1 expression in sensory neurons and CD4^+^T cells. (A) Time course of nociceptive thresholds in the EAP model in wild-type (WT, PD-1^fl/fl^), CD4^Cre^;PD-1^fl/fl^, Advillin^Cre^;PD-1^fl/fl^, and CD4^Cre^; Advillin^Cre^;PD-1^fl/fl^ mice. All genotypes exhibited progressive EAP-associated changes prior to treatment. SELTA administration (indicated by arrow) induced a reduction in pelvic hypersensitivity over the next 7 days compared to pretreatment baseline. In contrast, mice lacking PD-1 in sensory neurons, CD4^+^ T cells, or both showed no reduction following SELTA treatment. (B) Quantification of the treatment response showing SELTA-induced change in nociceptive threshold for each genotype, calculated as the difference between post-treatment and pre-treatment baseline values (Δ post-SELTA − pre-SELTA). WT mice demonstrated a significant reduction in responses while all conditional PD-1 knockout groups exhibited no reduction and showed progressive increase in responses over the 7-day period. No additional difference was observed in the double knockout compared with either single knockout, indicating a non-redundant requirement for PD-1 expression in both compartments. Data are shown as mean ± SEM. Statistical analysis was performed comparing the day 7 posttreatment response of each group to the corresponding WT using two-way ANOVA with Tukey’s multiple-comparisons test; \**P* < 0.05; **P < 0.01; \*\*\*\**P* < 0.0001

To isolate the SELTA-specific treatment effect and account for baseline differences across genotypes, we quantified the change in nociceptive threshold for each mouse relative to its pre-treatment baseline (Δ post-SELTA − pre-SELTA; Fig. 6B). Wild-type mice demonstrated a significant positive Δ response following SELTA treatment, whereas all conditional PD-1 knockout groups exhibited significantly blunted Δ responses. No additional reduction was observed in the double knockout compared with either single knockout. Together, these data demonstrate that PD-1 expression in both sensory neurons and CD4^+^ T cells is required for the full analgesic effect of SELTA in the EAP model.

## Discussion

This study identifies a neuroimmune mechanism in which a commensal-derived LTA engages inflammation-dependent PD-1 signaling to suppress pelvic pain. As summarized in the model (Fig. 7) SELTA activates TLR2/6-dependent NF-κB signaling, increases Pdcd1 transcription and PD-1^Y248^ phosphorylation in DRG neurons under inflammatory conditions, suppresses ATP-evoked calcium responses, and reduces pelvic hypersensitivity in EAP.

**Figure 7.**
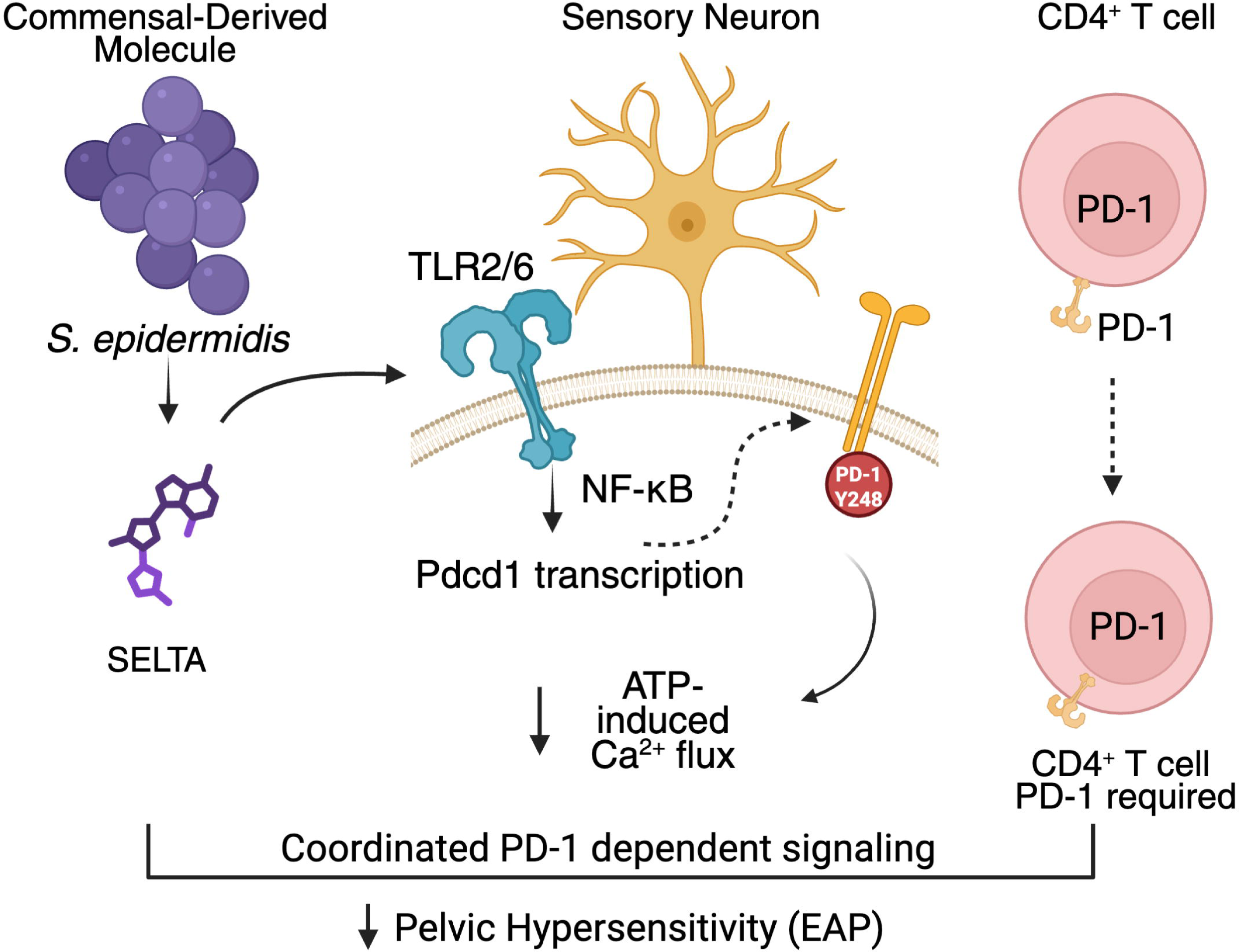
Proposed model of SELTA-mediated PD-1–dependent modulation of pelvic hypersensitivity. SELTA, a lipoteichoic acid (LTA) derived from a commensal *Staphylococcus epidermidis* strain (left), engages TLR2/6 heterodimers on sensory neurons and activates NF-κB–associated signaling. TLR2/6 activation is associated with increased *Pdcd1* transcription and enhanced PD-1 receptor expression in dorsal root ganglion (DRG) neurons. SELTA also increases phosphorylation of PD-1 at tyrosine 248 (Y248) within its intracellular domain. PD-1 signaling is associated with reduced ATP-induced Ca^2+^ influx in DRG neurons. In parallel, PD-1 expression in CD4⁺ T cells is required for the full analgesic effect of SELTA in vivo. Conditional deletion of PD-1 in sensory neurons or CD4⁺ T cells attenuate SELTA-mediated analgesia. These pathways converge to reduce pelvic hypersensitivity in experimental autoimmune prostatitis (EAP). Solid arrows indicate experimentally supported relationships; dashed arrows denote inferred or indirect associations.

PD-1 is increasingly recognized as an inhibitory receptor in sensory neurons. Engagement of PD-1 by PD-L1 suppresses nociceptor firing and inflammatory pain [6], whereas PD-1 blockade enhances pain sensitivity and interferes with opioid antinociception [8; 33]. Our findings extend this paradigm by demonstrating that a commensal-derived molecule can induce PD-1 expression and phosphorylation in sensory neurons during inflammation. Baseline single-cell transcriptomic analyses report minimal Pdcd1 expression in naïve DRG neurons [30], consistent with our observation that robust PD-1 upregulation occurs primarily under inflammatory stimulation. These findings support a model in which neuronal PD-1 functions as an inducible checkpoint engaged during peripheral sensitization rather than as a constitutive regulator of excitability.

Mechanistically, SELTA selectively activated TLR2/6 heterodimers, consistent with canonical LTA recognition [27]. While TLR signaling is often associated with pro-inflammatory programs, downstream transcriptional outputs can be context dependent. In inflamed sensory neurons, TLR2/6 activation was associated with increased Pdcd1 transcription and PD-1 phosphorylation. Although direct promoter-level interactions were not examined, these data place PD-1 downstream of TLR2/6 signaling in this system.

Importantly, SELTA-induced analgesia required PD-1 expression in both sensory neurons and CD4⁺ T cells. Although Advillin^Cre^ predominantly targets primary sensory neurons, limited off-target expression has been reported [11]. Conditional deletion of Pdcd1 in advillin-expressing neurons or CD4⁺ T cells significantly attenuated SELTA efficacy, and combined deletion did not further diminish the response. Baseline nociceptive behavior and EAP development were preserved across genotypes, indicating that PD-1 is not required for pain induction but is necessary for SELTA-mediated analgesia. These findings support a requirement for PD-1 expression in both neuronal and immune compartments for the full analgesic effect of SELTA.

An important consideration is the strain specificity of SELTA’s analgesic effect. While SELTA was purified from a prostate-localized *S. epidermidis* strain with anti-nociceptive properties [17], not all prostate-derived *S. epidermidis* isolates share this phenotype. In prior work from our laboratory, a different prostate-derived strain failed to attenuate pelvic pain and instead promoted inflammatory responses and hypersensitivity [15]. These findings underscore functional heterogeneity among *S. epidermidis* strains and suggest that structural differences in bacterial components, including lipoteichoic acids, critically determine host signaling outcomes. LTA structural features are known to influence TLR2 heterodimer engagement and downstream signaling bias [27]. Thus, the divergent pain phenotypes observed between isolates likely reflect discrete molecular characteristics rather than generalized species-level effects.

In summary, these findings describe a pathway linking a commensal-derived molecule to TLR2/6 activation, PD-1 signaling, and modulation of inflammatory pain. Further work will be required to define the molecular intermediates and to determine how neuronal and immune PD-1 signaling interact in this context.

## Declarations

### Ethics approval and consent to participate

All animal experiments were conducted in accordance with institutional guidelines and were approved by the Northwestern University Institutional Animal Care and Use Committee (IACUC).

### Consent for publication

Not applicable.

### Availability of data and materials

The datasets generated and/or analyzed during the current study are available from the corresponding author on reasonable request.

### Authors’ contributions

ZL performed experiments, acquired and analyzed data, and drafted the manuscript. YHC assisted with neuronal experiments and data analysis. CVO assisted with experimental work and data processing. MM contributed to data interpretation and provided expertise in neuronal physiology. AJS contributed to study design and interpretation. PT conceived and directed the study, supervised all aspects of the work, analyzed data, and wrote the manuscript. All authors read and approved the final manuscript.

## Acknowledgements

The authors thank Northwestern University core facilities for technical support.

## Authors’ information

PT is a Professor in the Department of Urology at Northwestern University Feinberg School of Medicine with a research focus on neuroimmune mechanisms in chronic pelvic pain.

## Data Availability

All data generated in this study are available from the corresponding author upon request.

---

**Supplementary Figure S1. Functional validation of HEK-hTLR2/1 and HEK-hTLR2/6 reporter cell lines.** HEK-Blue reporter cells stably expressing human TLR2/1 or TLR2/6 were stimulated with increasing concentrations of canonical TLR2 heterodimer agonists, and NF-κB activation was quantified by SEAP reporter activity (OD 655 nm). (A) HEK-Blue-hTLR2/1 cells were stimulated with Pam3CSK4 (TLR2/1 agonist) or Pam2CSK4 (TLR2/6 agonist). Pam3CSK4 induced robust NF-κB activation in TLR2/1-expressing cells, whereas Pam2CSK4 produced minimal activation, confirming functional TLR2/1 specificity. (B) HEK-Blue-hTLR2/6 cells were stimulated with Pam2CSK4 or Pam3CSK4. Pam2CSK4 induced strong NF-κB activation in TLR2/6-expressing cells, while Pam3CSK4 elicited significantly lower responses, confirming selective TLR2/6 signaling. Data are presented as mean ± SEM from independent experiments (individual data points shown). Statistical analysis was performed using one-way ANOVA with multiple comparisons; significance is indicated as ****p < 0.0001, and ns denotes not significant. These data confirm the functional integrity and ligand selectivity of the engineered HEK reporter lines used for SELTA signaling studies.

**Supplementary Figure S2. Validation of anti–PD-1, anti–phospho-PD-1 (Y248), and anti-TLR2 antibodies in mouse dorsal root ganglia.** Representative confocal images of 10-µm DRG sections from adult C57BL/6 mice stained for total PD-1, phospho-PD-1 (Y248), or TLR2 (green) together with the neuronal marker NeuN (red) (upper panels). Immunoreactivity is observed in NeuN-positive sensory neurons. To assess specificity, adjacent serial sections were processed in parallel with omission of the respective primary antibodies while retaining anti-NeuN and identical fluorophore-conjugated secondary antibodies (lower panels). No green signal was detected in no-primary controls, indicating minimal nonspecific secondary antibody binding. Additional peptide competition controls for total PD-1 are shown in Supplementary Figure S5. Antibody sources and catalog numbers are provided in the Methods. All images were acquired using identical imaging settings across experimental and control sections. Scale bar, 50 µm.

**Supplementary Figure S3. Single and repeated dosing of SELTA in WT EAP mice.** (A) Response frequency graph demonstrating pelvic tactile allodynia in control mice over 119 days of EAP. (B) Treatment of EAP with saline at days 21 and 70 followed by measurement of response frequency for each filament force. (C) Treatment with SELTA at days 21 and 70 and measurement of response frequency over time.

**Supplementary Figure S4. Concentration-dependent efficacy of SELTA in EAP.** (A) Response frequency graphs of control EAP (A) and increasing concentrations of SELTA (B-E) in EAP mice. Mice were treated at day 21 and therapeutic efficacy was evaluated at days 22,24, 26 and 28. Response frequency for each filament force over time is plotted.

**Supplementary Figure S5. Generation of the PD-1 conditional knockout (CKO) mouse and confirmation of PD-1 knockdown in sensory neurons.** (A) The breeding schematic for generation of sensory-specific PD-1 CKO mice involved crossing floxed PD-1 and somatic sensory neuron-specific advillin promotor-controlled Cre mice. (B) Confirmation of PD-1 knockdown in advillin-positive peripheral sensory neurons was done with PD-1 immunofluorescent staining. DRG from PD-1^fl/fl^ mice was incubated with anti-PD-1 antibody with or without a PD-1 blocking peptide to establish specificity of PD-1 staining. Knockdown of PD-1 expression was then confirmed in Advil^Cre^-PD-1^fl/fl^ mouse DRG Specificity of the knockdown to sensory neurons was verified by showing reduced expression in NeuN expressing cells but not in IBA-1-positive cells (white arrowheads). Scale bar=25 µm.

**Supplementary Figure S6. Validation of CD4^+^T cell–specific Pdcd1 deletion.** (A, C) Representative PCR genotyping demonstrating detection of the CD4^Cre^ transgene and floxed Pdcd1 alleles. (B) Immunofluorescence staining of lymph node sections showing loss of PD-1 immunoreactivity (red) in CD4⁺ T cell regions of PD-1^fl/fl^;CD4^Cre+/−^ mice compared with PD-1^fl/fl^ controls. Insets show higher magnification of boxed regions. (D) Flow cytometric analysis of splenocytes confirming reduced PD-1 expression in CD4⁺ T cells from CD4^Cre^ conditional knockout mice relative to wild-type controls.

**Supplementary Figure S7. Baseline behavioral characterization of PD-1 sensory neuron–specific conditional knockout mice.** Baseline behavioral assays were performed in 5–7-week-old PD-1^fl/fl^ (equal to WT) and PD-1^f/f^;Advillin^Cre^ (CKO) mice (n = 9 per genotype) to confirm that sensory-neuron PD-1 deletion does not alter general behavior. (A) Dark–light box test showing similar time spent in the light compartment in WT and CKO mice. (B) Y-maze spontaneous alternation showing comparable working memory and exploratory behavior across genotypes. (C) Novel object recognition test demonstrating similar discrimination indices in WT and CKO mice. (D) Open field test showing comparable center-time percentages, indicating no differences in baseline locomotor or anxiety-related behavior.

**Supplementary Figure S8. PD-1 expression in neurons is required for SELTA-induced attenuation of pelvic allodynia in EAP mice.** (A) Response frequency graph demonstrating pelvic tactile allodynia in EAP-treated PD-1^f/f^ mice. EAP was allowed to develop for 50 days before mice received a single intraurethral SELTA instillation. Pelvic tactile hypersensitivity was then assessed at 1, 3, 5, and 7 days after SELTA treatment using a graded series of von Frey filaments. Data are expressed as percentage increase in response frequency relative to baseline for each filament force. (B) Pelvic tactile allodynia in EAP-treated PD-1^f/f^;Advillin^Cre^ mice subjected to the same protocol. EAP was allowed to progress for 50 days, followed by intraurethral SELTA instillation and allodynia assessment at 1-, 3-, 5-, and 7-days post-treatment.

**Supplementary Figure S9. PD-1 expression in both neurons and CD4 T cells is required for SELTA-induced attenuation of pelvic allodynia in EAP mice.** (A) Response frequency graph demonstrating pelvic tactile allodynia in EAP-treated PD-1^fl/fl^ mice. EAP was allowed to develop for 21 days before mice received a single intraurethral SELTA instillation. Pelvic tactile hypersensitivity was then assessed at 1, 3, 5, and 7 days after SELTA treatment using a graded series of von Frey filaments. Data are expressed as percentage increase in response frequency relative to baseline for each filament force. Pelvic tactile allodynia was assessed in EAP-treated Advillin^Cre^;PD-1^fl/fl^ (B) CD4^Cre^;PD-1^fl/fl^ (C) and CD4^Cre^;Advillin^Cre^;PD-1^fl/fl^ (D)mice.

