## Supplementary Figures S1-S9 for "A Commensal-Derived Lipoteichoic Acid Engages an Inducible Neuronal PD-1 Checkpoint to Suppress Inflammatory Pain"

Supplementary Figure S1

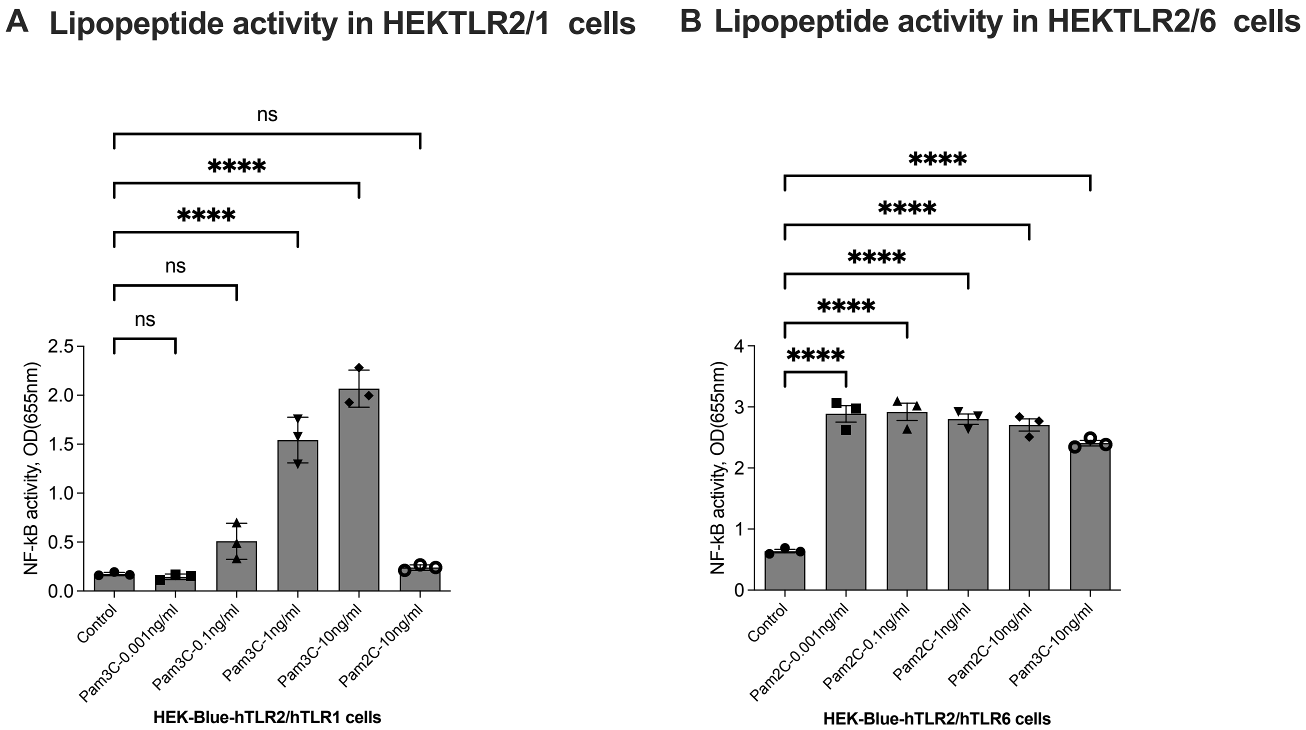


**Supplementary Figure 1. Functional validation of HEK-hTLR2/1 and HEK-hTLR2/6 reporter cell lines.** HEK-Blue reporter cells stably expressing human TLR2/1 or TLR2/6 were stimulated with increasing concentrations of canonical TLR2 heterodimer agonists, and NF-κB activation was quantified by SEAP reporter activity (OD 655 nm). (A) HEK-Blue-hTLR2/1 cells were stimulated with Pam3CSK4 (TLR2/1 agonist) or Pam2CSK4 (TLR2/6 agonist). Pam3CSK4 induced robust NF-κB activation in TLR2/1-expressing cells, whereas Pam2CSK4 produced minimal activation, confirming functional TLR2/1 specificity. (B) HEK-Blue-hTLR2/6 cells were stimulated with Pam2CSK4 or Pam3CSK4. Pam2CSK4 induced strong NF-κB activation in TLR2/6-expressing cells, while Pam3CSK4 elicited significantly lower responses, confirming selective TLR2/6 signaling. Data are presented as mean ± SEM from independent experiments (individual data points shown). Statistical analysis was performed using one-way ANOVA with multiple comparisons; significance is indicated as ****$p < 0.0001$, and ns denotes not significant. These data confirm the functional integrity and ligand selectivity of the engineered HEK reporter lines used for SELTA signaling studies.

#### Supplementary Figure S2

##### Validation for anti-PD-1, anti-phospho-PD-1 (Y248) and anti-TLR2 antibodies

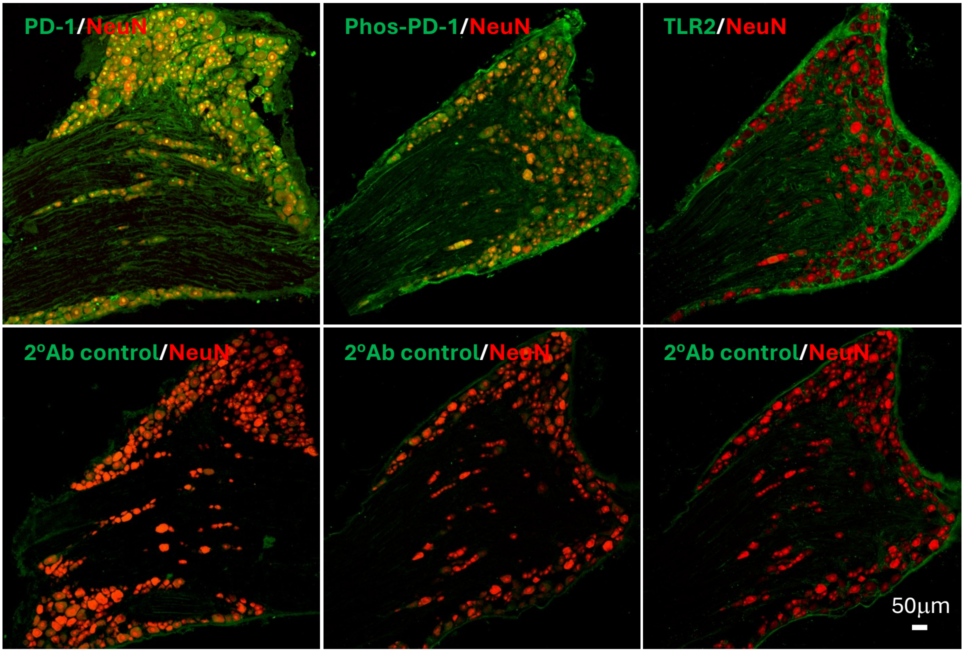


**Supplementary Figure S2. Validation of anti–PD-1, anti–phospho-PD-1 (Y248), and anti-TLR2 antibodies in mouse dorsal root ganglia.** Representative confocal images of 10-μm DRG sections from adult C57BL/6 mice stained for total PD-1, phospho-PD-1 (Y248), or TLR2 (green) together with the neuronal marker NeuN (red) (upper panels). Immunoreactivity is observed in NeuN-positive sensory neurons. To assess specificity, adjacent serial sections were processed in parallel with omission of the respective primary antibodies while retaining anti-NeuN and identical fluorophore-conjugated secondary antibodies (lower panels). No green signal was detected in no-primary controls, indicating minimal nonspecific secondary antibody binding. Additional peptide competition controls for total PD-1 are shown in Supplementary Figure S3. Antibody sources and catalog numbers are provided in the Methods. All images were acquired using identical imaging settings across experimental and control sections. Scale bar, 50 μm.

Supplementary Figure S3

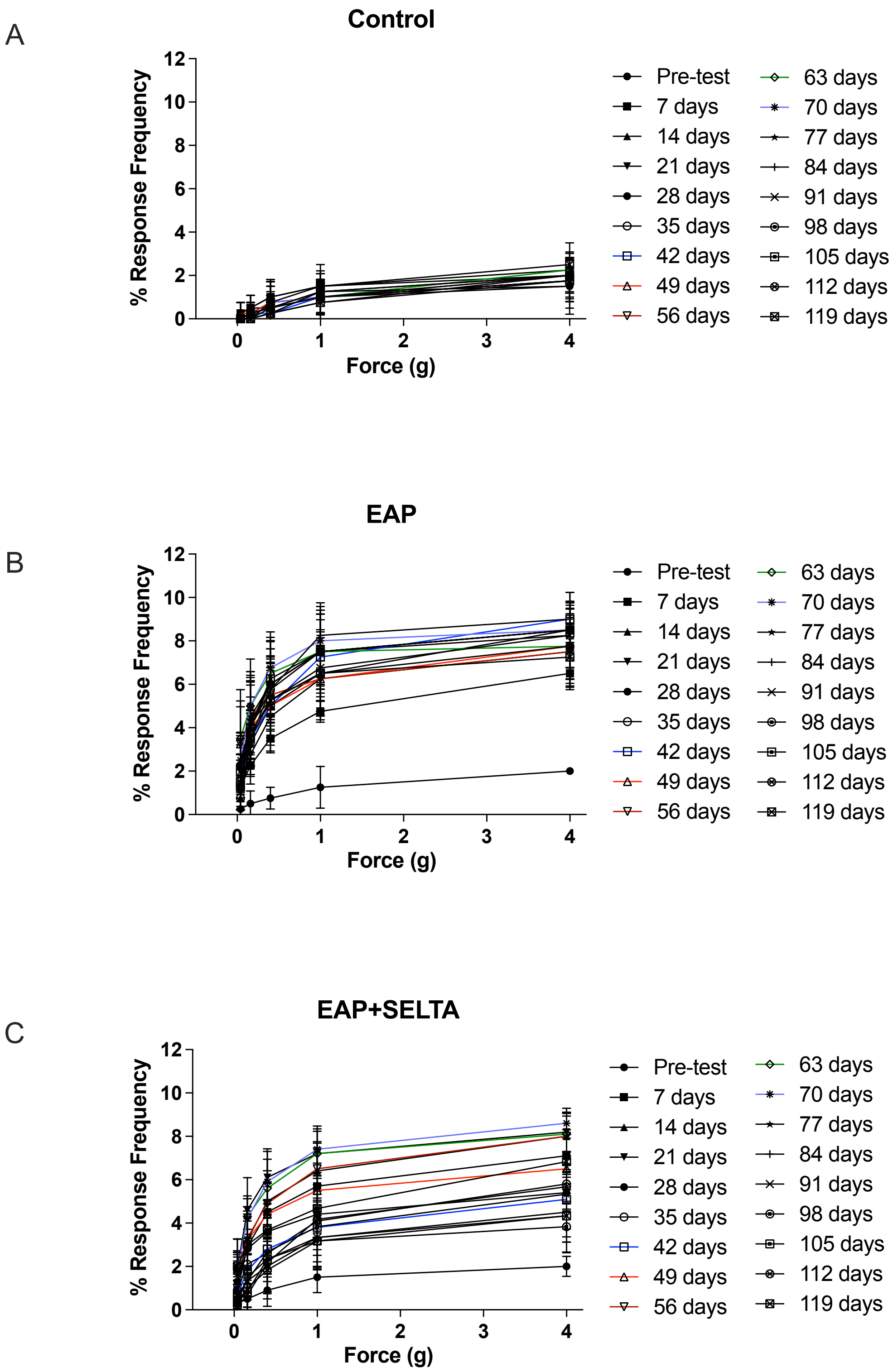


**Supplementary Figure S3. Single and repeated dosing of SELTA in WT EAP mice..** (A) Response frequency graph demonstrating pelvic tactile allodynia in control mice over 119 days of EAP. (B) Treatment of EAP with saline at days 21 and 70 followed by measurement of response frequency for each filament force. (C) Treatment with SELTA at days 21 and 70 and measurement of response frequency over time.

#### Supplementary Figure S4

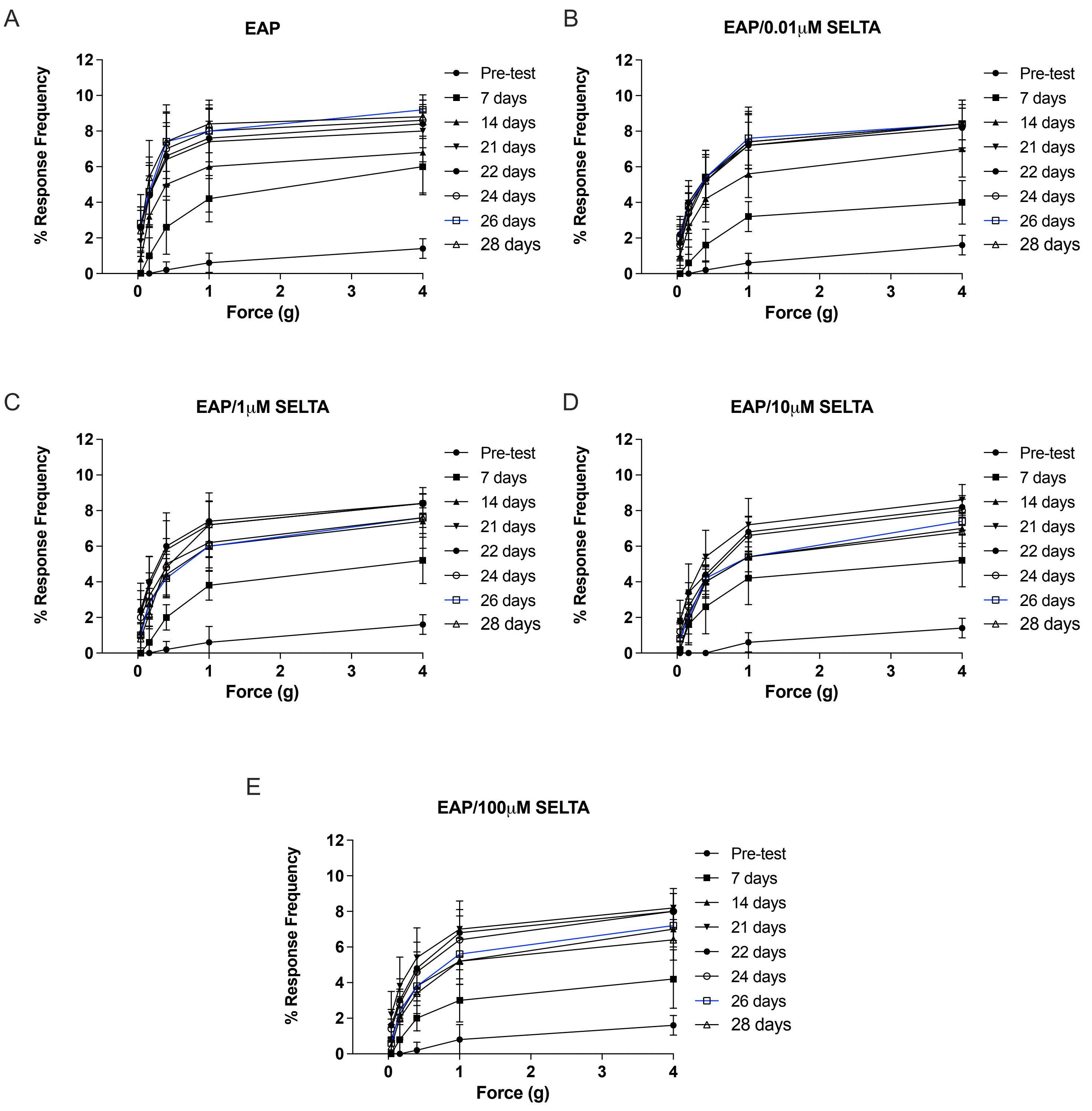


**Supplementary Figure S4. Concentration-dependent efficacy of SELTA in EAP.** (A) Response frequency graphs of control EAP (A) and increasing concentrations of SELTA (B-E) in EAP mice. Mice were treated at day 21 and therapeutic efficacy was evaluated at days 22,24, 26 and 28. Response frequency for each filament force over time is plotted.

#### Supplementary Figure S5

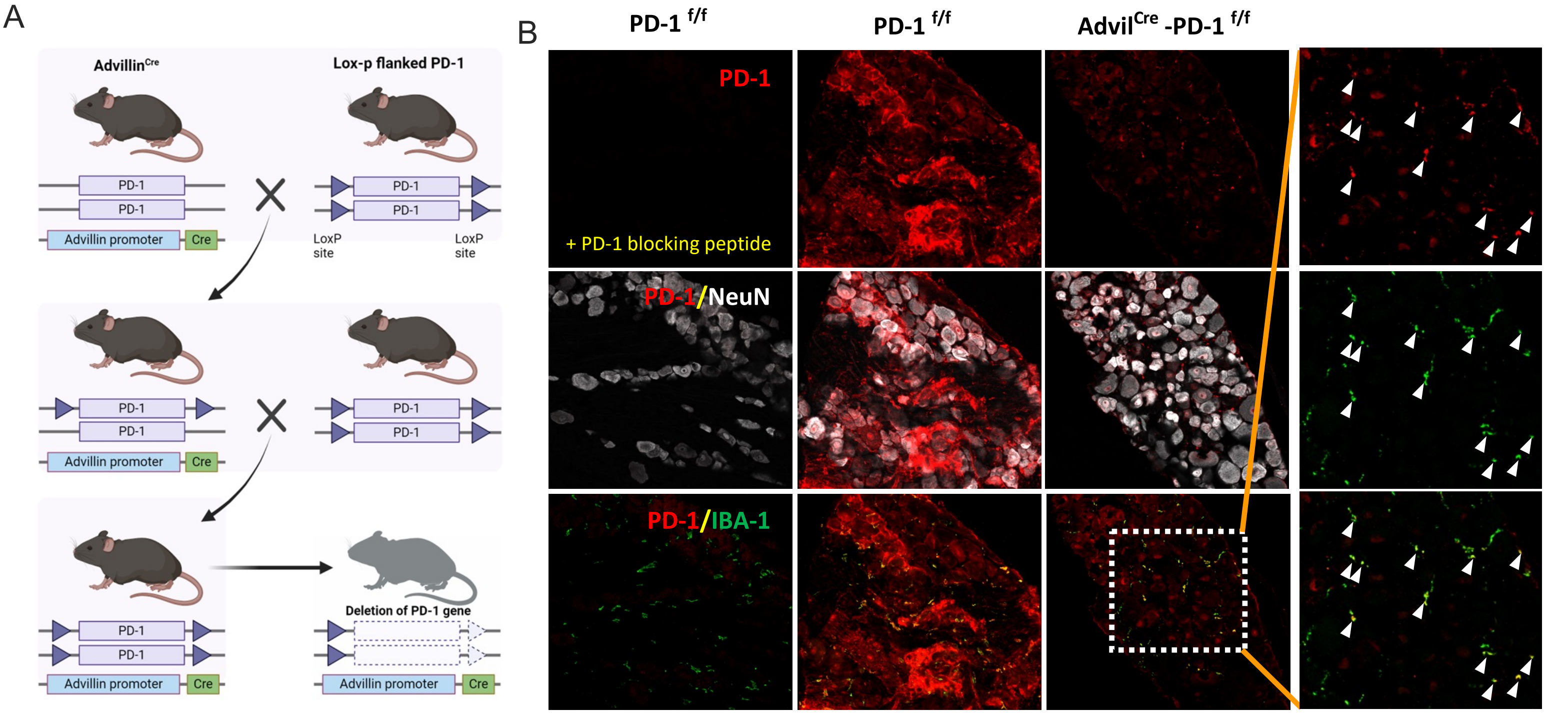


**Supplementary Figure S5. Generation of the PD-1 conditional knockout (CKO) mouse and confirmation of PD-1 knockdown in sensory neurons.** (A) The breeding schematic for generation of sensory-specific PD-1 CKO mice involved crossing floxed PD-1 and somatic sensory neuron-specific advillin promotor-controlled Cre mice. (B) Confirmation of PD-1 knockdown advillin-positive periphery sensory neurons was the CKO was done with PD-1 immunofluorescent staining. DRG from PD-1$^{f/f}$ mice was incubated with anti-PD-1 antibody with or without a PD-1 blocking peptide to establish specificity of PD-1 staining. Knockdown of PD-1 expression was then confirmed in Advil$^{cre}$-PD-1$^{f/f}$ mouse DRG. Specificity of the knockdown to sensory neurons was verified by showing reduced expression in NeuN expressing cells but not in IBA-1-positive cells (white arrowheads). Scale bar=25 μm.

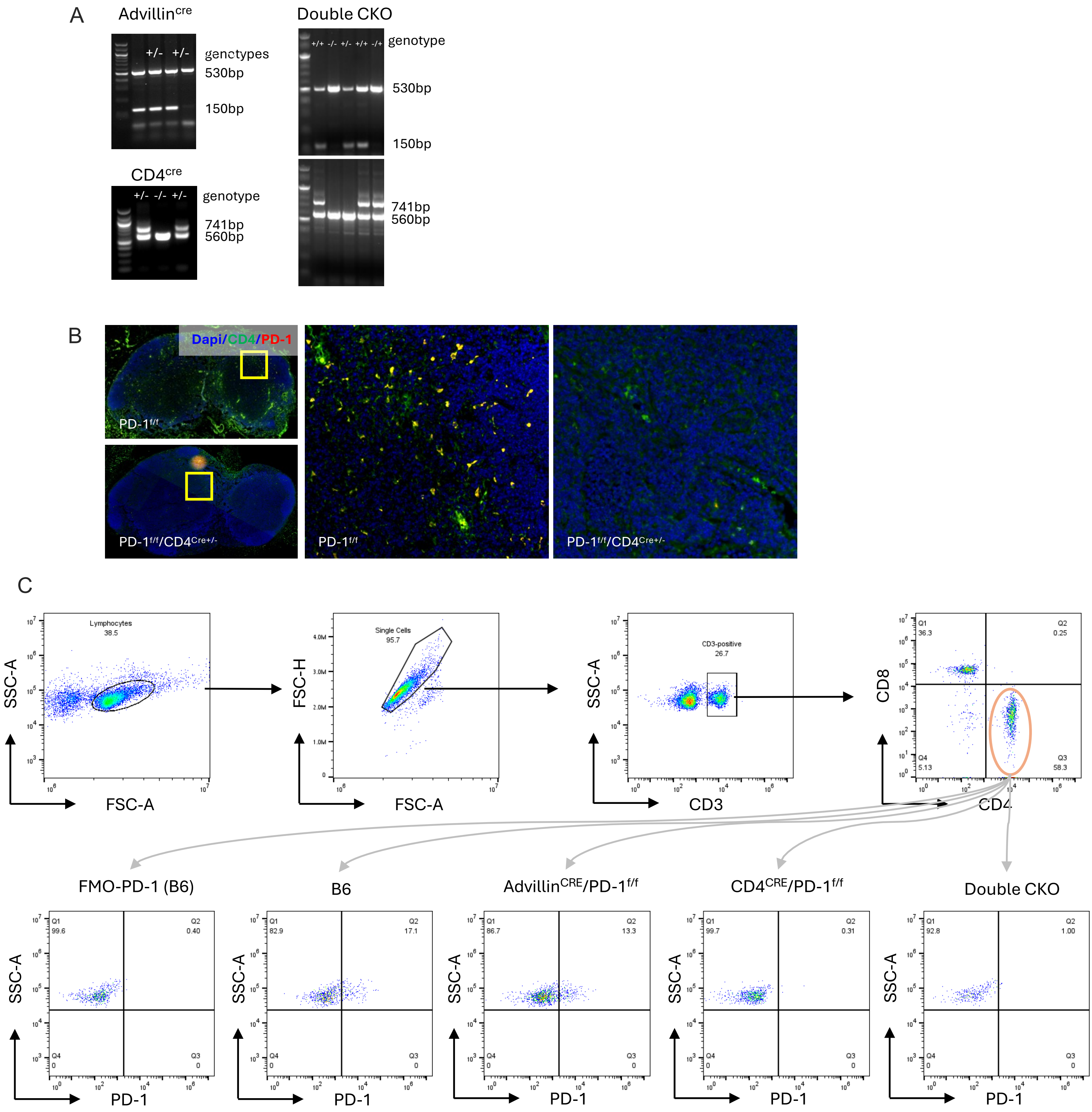


**Supplementary Figure S6. Validation of Pdcd1 deletion and reduction in CD4 PD-1 expression in PD-1$^{f/f}$;CD4-Cre$^{+/-}$, and Double CKO mice.**

(A, C) Representative PCR genotyping from all CKO mice. (B) Immunofluorescence staining of lymph node sections showing loss of PD-1 immunoreactivity (red) in CD4$^{+}$ T cell regions of Pdcd1^f/f^;CD4-Cre^+/–^ mice compared with Pdcd1^f/f^ controls. Insets show higher magnification of boxed regions. (C) Flow cytometric analysis of PD-1 expression in splenocytes from Pdcd1^f/f^, Pdcd1^f/f^;Advillin-Cre^+/–^, Pdcd1^f/f^;CD4-Cre^+/–^, and double CKO mice.

Supplementary Figure S7

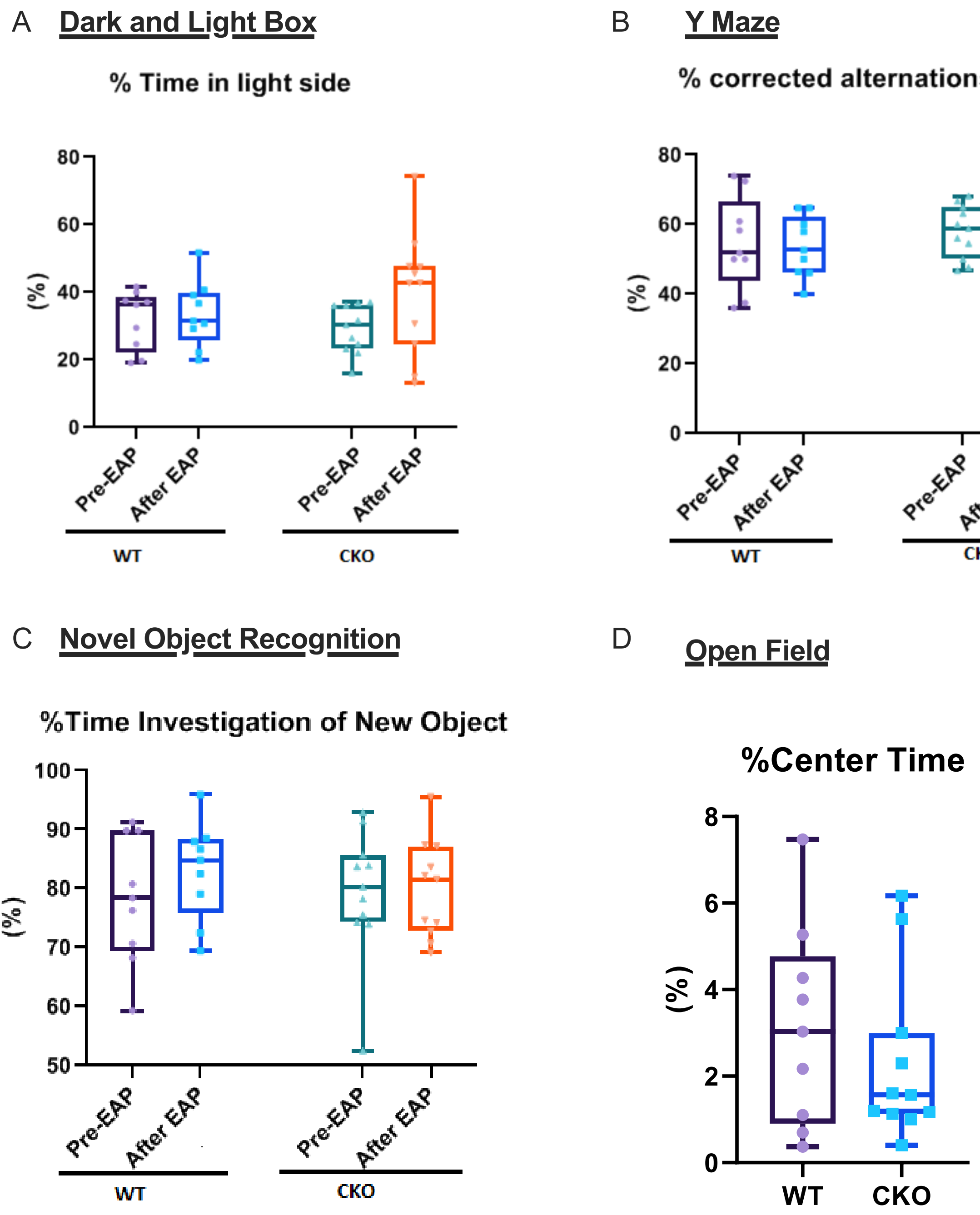


**Supplementary Figure S7. Baseline behavioral characterization of PD-1 sensory neuron–specific conditional knockout mice.** Baseline behavioral assays were performed in 5–7-week-old PD-1$^{f/f}$ (WT) and PD-1$^{f/f}$;Advillin-Cre (CKO) mice (n = 9 per genotype) to confirm that sensory-neuron PD-1 deletion does not alter general behavior. (A) Dark–light box test showing similar time spent in the light compartment in WT and CKO mice. (B) Y-maze spontaneous alternation showing comparable working memory and exploratory behavior across genotypes. (C) Novel object recognition test demonstrating similar discrimination indices in WT and CKO mice. (D) Open field test showing comparable center-time percentages, indicating no differences in baseline locomotor or anxiety-related behavior.

Supplementary Figure S8

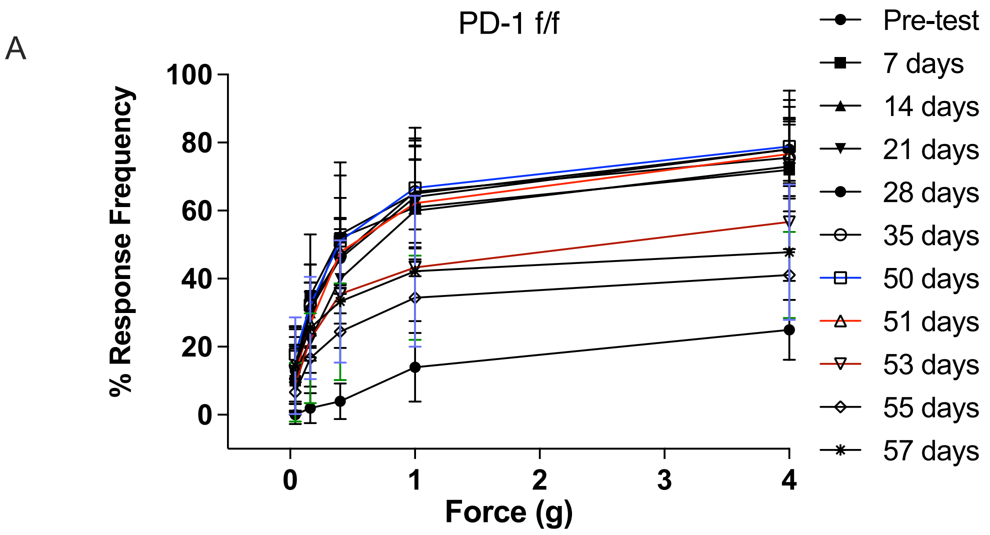


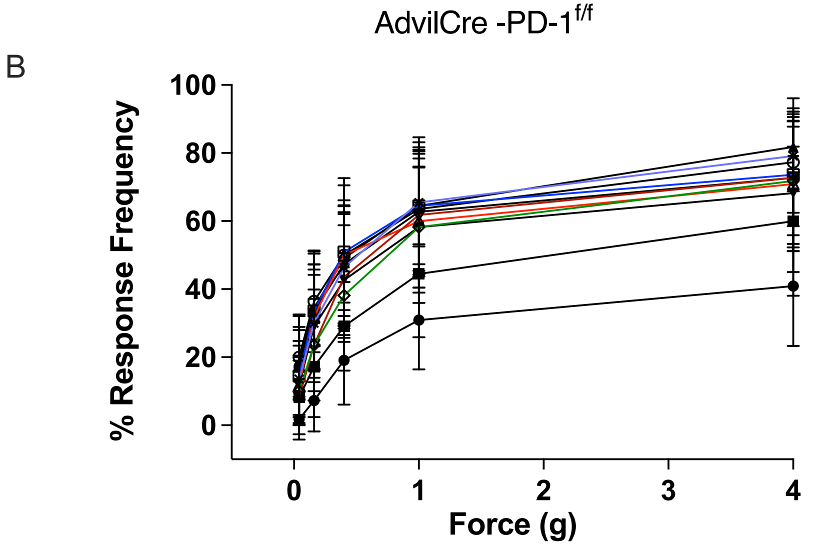


**Supplementary Figure S8. PD-1 expression in neurons is required for SELTA-induced attenuation of pelvic allodynia in EAP mice.** (A) Response frequency graph demonstrating pelvic tactile allodynia in EAP-treated PD-1$^{f/f}$ mice. EAP was allowed to develop for 50 days before mice received a single intraurethral SELTA instillation. Pelvic tactile hypersensitivity was then assessed at 1, 3, 5, and 7 days after SELTA treatment using a graded series of von Frey filaments. Data are expressed as percentage increase in response frequency relative to baseline for each filament force. (B) Pelvic tactile allodynia in EAP-treated PD-1$^{f/f}$;Advillin-Cre mice subjected to the same protocol. EAP was allowed to progress for 50 days, followed by intraurethral SELTA instillation and allodynia assessment at 1-, 3-, 5-, and 7-days post-treatment.

### Supplementary Figure S9

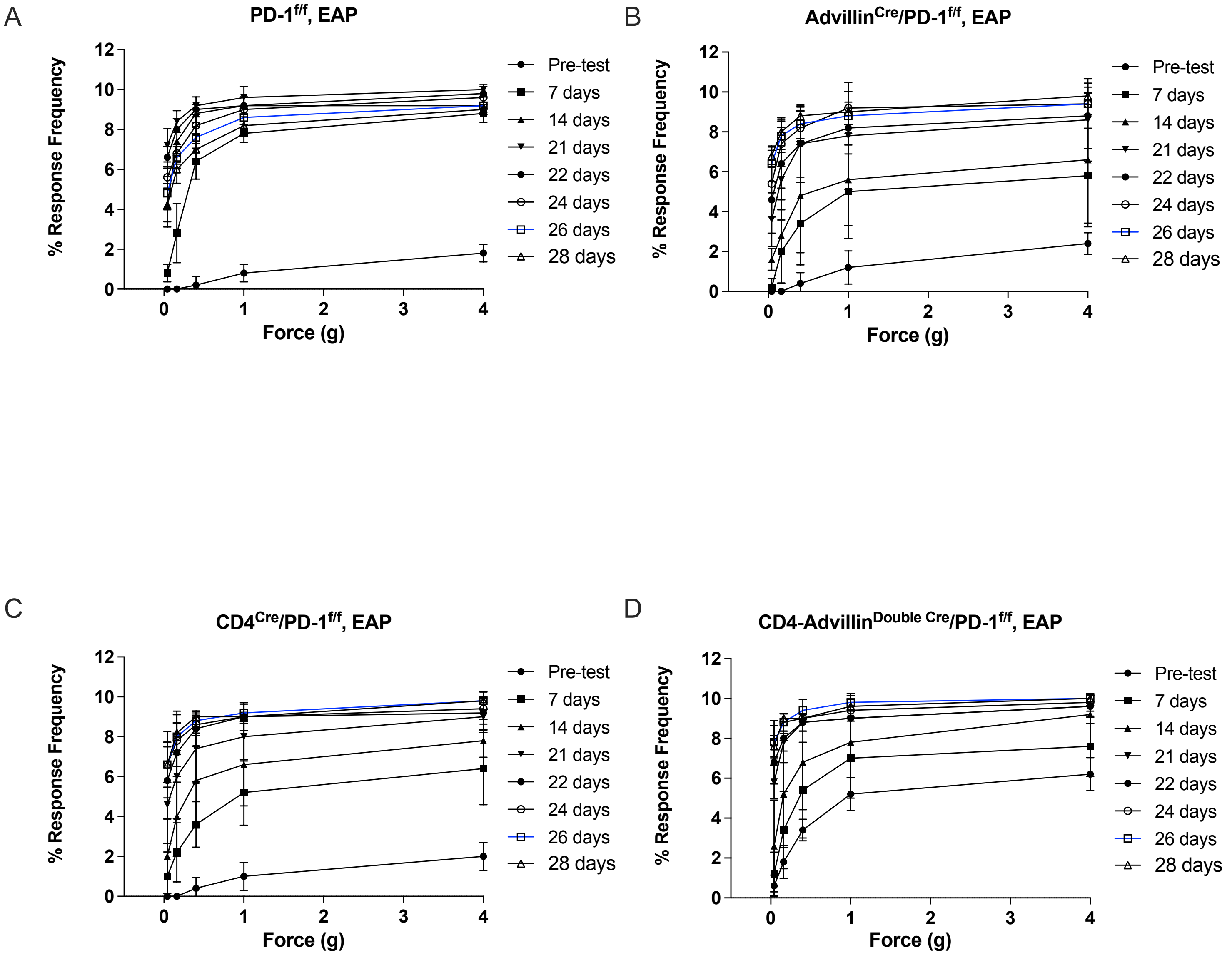


**Supplementary Figure S9. PD-1 expression in both neurons and CD4$^+$ T cells is required for SELTA-induced attenuation of pelvic allodynia in EAP mice.** (A) Response frequency graph demonstrating pelvic tactile allodynia in EAP-treated PD-1$^{f/f}$ mice. EAP was allowed to develop for 21 days before mice received a single intraurethral SELTA instillation. Pelvic tactile hypersensitivity was then assessed at 1, 3, 5, and 7 days after SELTA treatment using a graded series of von Frey filaments. Data are expressed as percentage increase in response frequency relative to baseline for each filament force. Pelvic tactile allodynia was assessed in EAP-treated Advillin-Cre;PD-1$^{fl/fl}$(B) CD4-Cre;PD-1$^{fl/fl}$ (C) and CD4-Cre;Advillin-Cre;PD-1$^{fl/fl}$ (D) mice.
